# Dimensional and Categorical Solutions to Parsing Depression Heterogeneity in a Large Single-Site Sample

**DOI:** 10.1101/2023.07.05.547873

**Authors:** Katharine Dunlop, Logan Grosenick, Jonathan Downar, Fidel Vila-Rodriguez, Faith M. Gunning, Zafiris J. Daskalakis, Daniel M. Blumberger, Conor Liston

## Abstract

**Background:** Recent studies have reported significant advances in modeling the biological basis of heterogeneity in major depressive disorder (MDD), but investigators have also identified important technical challenges, including scanner-related artifacts, a propensity for multivariate models to overfit, and a need for larger samples with deeper clinical phenotyping. The goals of this work were to develop and evaluate dimensional and categorical solutions to parsing heterogeneity in depression that are stable and generalizable in a large, deeply phenotyped, single-site sample.

**Methods:** We used regularized canonical correlation analysis (RCCA) to identify data-driven brain-behavior dimensions explaining individual differences in depression symptom domains in a large, single-site dataset comprising clinical assessments and resting state fMRI data for N=328 patients with MDD and N=461 healthy controls. We examined the stability of clinical loadings and model performance in held-out data. Finally, hierarchical clustering on these dimensions was used to identify categorical depression subtypes

**Results:** The optimal RCCA model yielded three robust and generalizable brain-behavior dimensions explaining individual differences in depressed mood and anxiety, anhedonia, and insomnia. Hierarchical clustering identified four depression subtypes, each with distinct clinical symptom profiles, abnormal RSFC patterns, and antidepressant responsiveness to repetitive transcranial magnetic stimulation.

**Conclusions:** Our results define dimensional and categorical solutions to parsing neurobiological heterogeneity in MDD that are stable, generalizable, and capable of predicting treatment outcomes, each with distinct advantages in different contexts. They also provide additional evidence that RCCA and hierarchical clustering are effective tools for investigating associations between functional connectivity and clinical symptoms.

## INTRODUCTION

Major depressive disorder (MDD) is a heterogeneous syndrome widely assumed to comprise multiple distinct subtypes associated with differing pathophysiological processes and potentially requiring different treatments, but there is no consensus on how to define them. MDD is diagnosed when a patient presents with five or more of nine symptom criteria, including depressed mood, anhedonia, psychomotor symptoms, and changes in sleep or appetite, among others (1–3). Although anxiety is not a core diagnostic criterion, patients with MDD often present with comorbid anxiety disorders or subclinical anxiety symptoms. Understanding the biological basis of individual differences and heterogeneity in MDD is a critical challenge for developing new approaches to diagnosis and treatment.

Pioneering early work on this topic sought to differentiate MDD subtypes based on patterns of co-occurring symptoms or other clinical characteristics, such as melancholic, atypical, and seasonal depression (4–7). While this approach to clinical subtyping has enabled important advances in our understanding of heterogeneity, it has been challenging to identify stable biological substrates of clinical depression subtypes and to discover biomarkers capable of diagnosing depression or predicting treatment outcomes at the individual level (4,8–10). An alternative to this approach involves subtyping patients based on shared biological features (11–13), which are commonly defined using fMRI and EEG measures of functional connectivity in depression-related brain networks (12,14–19). This approach has shown promise in identifying subtypes of psychotic and affective disorders across the lifespan (20–22) and could eventually be used to predict responses to antidepressant medications, psychotherapy, and brain stimulation (23–30).

Recent efforts to parse the neurobiological basis of heterogeneity in depression have taken either a dimensional approach (21,31,32)—using multivariate models to identify brain-behavior dimensions that explain individual differences in depressive symptoms along a continuous spectrum—or a categorical approach (17,19,33), clustering subjects into relatively homogeneous categorical subgroups based on resting state functional connectivity patterns and other shared biological measures. While categorical models may be especially useful for clinicians, who are often trained to diagnose disorders and select treatments based on categorical heuristics, dimensional approaches may be more useful for some research purposes, especially for modeling data that are continuously distributed and in contexts where biological measures predict a continuous range of clinical symptom profiles. We (2,34) and others (35,36) have taken a hybrid approach, integrating dimensional and categorical models of heterogeneity in depression (31,35) and extending this method to generate robust, generalizable models of heterogeneity in other neuropsychiatric syndromes such as autism spectrum disorder (37). In our previous work, we used canonical correlation analysis (CCA(38)) to identify two brain-behavior dimensions defined by co-occurring patterns of resting-state functional connectivity (RSFC) and depressive symptoms: one dimension was related to anhedonia and psychomotor slowing, and the second was associated with anxiety and insomnia (2). Clustering on these dimensions yielded four MDD subtypes defined by distinct patterns of abnormal RSFC that were associated with differing clinical symptom profiles and varying responsiveness to repetitive transcranial magnetic stimulation. Although two brain-behavior dimensions and a four-cluster solution were optimal in this work, one question left unanswered was whether additional dimensions and higher-order clustering solutions could be identified in larger samples.

More recently, investigators have also identified important technical challenges to generating multivariate models of heterogeneity using fMRI data and clinical symptom scores. First, the approach used in our previous work is prone to overfitting (39), and analyses in one recent report indicate that extraordinarily large sample sizes may be required for detecting reproducible brain-behavior relationships, at least in the context of some behavioral measures and analytical approaches (40). We subsequently showed that overfitting could be mitigated by using regularization and a stabilized bootstrapped feature selection procedure (34), and others have shown that with appropriate methodology (e.g., regularization, cross-validation in held-out data), multivariate analyses can identify replicable effects with substantially smaller sample sizes (41). Second, scanner-related artifacts are substantial in fMRI data and may distort brain-behavior associations or render them undetectable; analyses in a larger dataset derived from a single site would eliminate these confounds (34,42,43). Third, deeper clinical phenotyping within a single-scanner sample may be useful for revealing brain-behavior associations that would not otherwise be detectable. For example, our previous work relied primarily on the Hamilton Depression Rating Scale (HRSD) for quantifying depressive symptoms (44). While this scale is widely used for assessing overall depressive severity and has many strengths, it was not designed to comprehensively assess specific symptom domains (45), including anhedonia—a core feature of depression that is evaluated based on a single item in the HAMD that conflates multiple dimensions of hedonic function with function at work and other domains (46).

Here, we set out to develop both dimensional and categorical solutions using updated statistical approaches to parsing heterogeneity in depression in a large, deeply phenotyped, single-site sample, comprising extensive clinical assessments and resting state fMRI data for N=328 patients with MDD. First, we used regularized CCA to identify robust and generalizable brain-behavior dimensions for understanding the neurobiological basis of individual differences in depression. Next, we used hierarchical clustering on these dimensions to identify categorical MDD subtypes, and we evaluated their stability and reproducibility. Finally, we characterized both models—dimensional and categorical—by examining associations between specific atypical RSFC features, clinical symptom profiles, and antidepressant responses to rTMS. The optimal RCCA model and clustering solution yielded three brain-behavior dimensions and four depression subtypes explaining individual differences in clinical symptoms, abnormal RSFC, and antidepressant responsiveness to repetitive transcranial magnetic stimulation.

## MATERIALS & METHODS

### Participants

Participants were recruited at Toronto Western Hospital and the Centre for Addiction and Mental in Toronto, Canada (2,47,48). All participants were enrolled in clinical studies assessing either the antidepressant efficacy of novel cortical targets for rTMS (i.e., dorsomedial prefrontal cortex [DMPFC] (2,48)) or were recruited a large non-inferiority trial testing the efficacy of novel rTMS protocols over the dorsolateral prefrontal cortex (DLPFC)(47). Inclusion/exclusion criteria and treatment designs are detailed in the supplemental methods; briefly, all participants were between 18-65 years old and met DSM-5 diagnostic criteria for major depressive disorder. We included data from individuals in this sample who were previously included in our original analysis, since the purpose of this new analysis was to expand upon our methods and evaluate performance under ideal conditions (i.e., using scans obtained from a single scanner). We also included fMRI data from 470 healthy, nondepressed participants, 96 of whom were acquired on the same scanner and with identical imaging parameters to the MDD sample (supplemental methods).

All participants provided written informed consent, and all studies were approved by the University Health Network, Centre for Addiction and Mental Health Research Ethics Boards or their respective Institutional Review Board.

### Clinical Measures

All MDD participants were assessed using the 17-item Hamilton Rating Scale for Depression(44) (HRSD). Item 17 of the HRSD (Insight) was omitted from all analyses because all participants had a score of zero on this item. Other clinical scales included the 21-item Beck Depression Inventory II(49) (BDI-II), and the 28-item Inventory of Depressive Symptomatology (50) (IDS). Because the BDI-II and IDS were not completed in all participants, we generated composite scores for five symptoms not fully captured by the HRSD, described in a subsequent section. Severity on the 16 HRSD items and five composite scores were used to generate brain-behavior dimensions. All measures were acquired at either the screening or baseline visit of their respective studies, and assessor-administered scales were acquired by trained evaluators.

### Composite Scores

All participants were assessed on the Hamilton Depression Rating Scale and either the Beck Depression Inventory (65.5%) or the Inventory of Depressive Symptoms (34.5%). To gain neurobiological insights on symptoms omitted from the HRSD, we included five composite scores using analogous items from the BDI-II and IDS: pessimism about the future (IDS#16, BDI-II#2); loss of interest/involvement (IDS#18, BDI-II#12); reduced pleasure/enjoyment (IDS#19, BDI-II#4); difficulty concentrating/decision-making (IDS#14, BDI-II#19 & 13); and irritability (IDS#8, BDI-II#17). Composite score generation is detailed in the supplemental methods.

### Neuroimaging Acquisition and Preprocessing

All MDD neuroimaging data were acquired at Toronto Western Hospital (Toronto, Canada) on a 3 Tesla GE Signa HDx equipped with an eight-channel phased-array head coil. The neuroimaging acquisition parameters have been described in detail elsewhere (51), and are outlined in the supplemental methods. Resting-state scans were preprocessed using the Analysis of Functional Neuroimages (AFNI) software package (52). Preprocessing was identical to that described in Drysdale et al.(2) (Supplemental Methods). Briefly, we censored a given volume and the volumes preceding and following it if head motion for that volume exceeded a threshold of 0.3 mm (Euclidean distance). 40 of 368 scanned MDD participants (10.9%) were excluded from further analysis because motion censoring rendered the number of volumes insufficient to complete denoising. Whole-brain RSFC matrices were extracted using the parcellation described in Drysdale et al. (2) (**Figure S1**), generating 33,153 functional connectivity features.

### CCA Optimization Overview

With the aim of assessing the generalizability and stability of CVs, we began by optimizing three hyperparameters for L2-norm regularized CCA (**Figure S2A**): the number of RSFC features, and the two regularization terms penalizing the RSFC and clinical matrices, λ1 and λ2, respectively. Each training iteration began by partitioning the data into a two-thirds (n=219) training and one-third outer fold (*n*=109; 100 replicates). We then further partitioned the outer fold training set into a two-thirds training (*n*=146) and one-third testing inner fold (*n*=73; 20 replicates). Within this inner training dataset, we performed bootstrapped feature selection to identify the most relevant feature. Next, we performed a grid search, modeling different combinations of CCA hyperparameters with the inner training set and assessing the canonical correlation in the inner test set. For each of the 100 outer replicates, the optimal hyperparameter combination was defined as the highest median canonical correlation in test data for CV1. We also assessed the impact of sample size, the strength of correlations, and clinical item range on CCA optimization, summarized in the Supplemental Methods and Results. Finally, we evaluated the optimal CCA parameters using the outer training set and projected to the outer test partition to generate a final canonical holdout canonical correlation.

### Dimension Significance Testing

With the optimal hyperparameters identified, we next sought to characterize generalizability and stability of CCA dimensions. We defined generalizability as how well each CCA dimension performed when applied unseen data, measured as the mean canonical correlation of RSFC and clinical dimension scores in held out data. We defined stability as the mean absolute similarity of clinical loadings for each dimension across all training replicates. To test the generalizability of CCA dimensions (**Figure S2B**), we began by partitioning the data into a two-thirds training and one-third test set (1,000 replicates). We performed bootstrapped feature selection using training data to identify the most relevant RSFC features and projected CCA model coefficients to the test dataset to evaluate the canonical correlation. We randomly permuted the testing data (1,000 replicates) to generate a null distribution of canonical correlations.

To test whether the clinical loadings for a canonical variate were significantly stable (**Figure S3C**), we randomly shuffled participants’ clinical data within the two-thirds training set (1,000 replicates). We also calculated the stability of clinical loadings, calculated as the average absolute Pearson’s correlation of clinical loadings across all 1000 replicates. For each iteration, we retained CV clinical loadings, generating a null distribution of clinical loading agreement across 1,000 null replicates.

We proceeded to evaluate subsequent CV generalizability using sequential orthogonal deflation, meaning that after we modeled a CV, we orthogonally deflated the data within each iteration to evaluate test canonical correlations for the subsequent CV. We corrected for multiple comparisons accounting for the fact that the canonical correlates are ranked, such that the family-wise error adjusted for the rank-*k* CV corresponded to the cumulative maximum p-value(53). We considered a CV to be significantly generalizable or stable at a corrected-α<0.05, one-tailed. We proceeded with assessing CVs until neither the generalizability nor the clinical loading stability was significant.

### Characterizing Significant CVs

We next generated a final regularized CCA model using RSFC and clinical data for all subjects for significant CVs, using tuned hyperparameters. We evaluated the Bonferroni-corrected clinical loadings and False-Discovery Rate-corrected (FDR) RSFC loadings for each CV.

### Hierarchical Clustering

To evaluate hierarchical clustering performance, we first extracted the Calinski-Harabasz Criterion(54) for 2-10 cluster solutions for combinations of significant CVs. We evaluated criterion values using normalized RSFC CV scores generated from the final model. The optimal cluster solution was identified as the maximum Calinski-Harabasz Criterion value. We used permutation testing (1,000 replicates) to evaluate hierarchical clustering significance. CV scores were generated using randomly permutated clinical and intact RSFC matrices, and Calinski-Harabasz scores were generated for potential solutions. Significant clustering solutions indicated a superior ratio of within to between cluster dispersion (Calinski-Harabasz Criterion) at *p*<0.05, one-tailed.

To test the stability of subtype membership, we generated cluster IDs in two-thirds subsamples of unshuffled data (1,000 replicates). CV scores were generated, and we evaluated the similarity of cluster membership from each subsample replicate to that achieved in the complete sample using adjusted mutual information. The optimal stability of cluster membership was defined as the elbow of the mean adjusted mutual information, using the Matlab command “knee_pt.” We tested cluster membership stability using the adjusted mutual information of two-thirds subsamples of shuffled clinical and intact RSFC data (1,000 iterations) relative to unshuffled cluster membership generated using the complete sample.

With the optimal number of clusters identified, we proceeded to characterize symptom and RSFC differences by subtype. First, we performed Kruskal-Wallis tests to identify significant differences in symptoms by subtype, and relative to symptom severity of the entire group. Next, we performed Wilcoxon rank sum tests for each RSFC feature and subtype to test for abnormal RSFC relative to healthy controls. Lastly, we identified differences in rTMS response and remission rates using chi-square tests and binomial logistic regression. Statistics were corrected for multiple comparisons using the FDR.

## RESULTS

### Three brain-behavior dimensions explaining individual differences in MDD

Our first aim was to define robust and reproducible brain-behavior dimensions explaining individual differences in distinct symptom domains in a deeply phenotyped sample of N=368 individuals with MDD, scanned and evaluated at a single site. Of the 368 participants recruited and scanned, 328 participants (215 female, mean age = 40.4 ± 12.1 [SD] years, range = 18-70; see Table 1 for demographic and clinical data) were retained following quality control (see Methods for details). Before attempting to build a multivariate model linking brain and behavior, we began by evaluating whether robust correlations between depressive symptoms and RSFC features were detectable in this sample, using bootstrapped feature selection to identify RSFC features that were stably correlated with a given clinical symptom score. We tested for significance using a permutation test to evaluate whether individual differences in a given clinical symptom were correlated with a number of RSFC features greater than that expected by chance. This analysis identified robust associations between clinical symptoms and RSFC features exceeding what would be expected by chance for 18 of 21 clinical symptoms (**Figure 1A**), with small to moderate effect sizes encompassing thousands of RSFC features.

**Figure 1:**
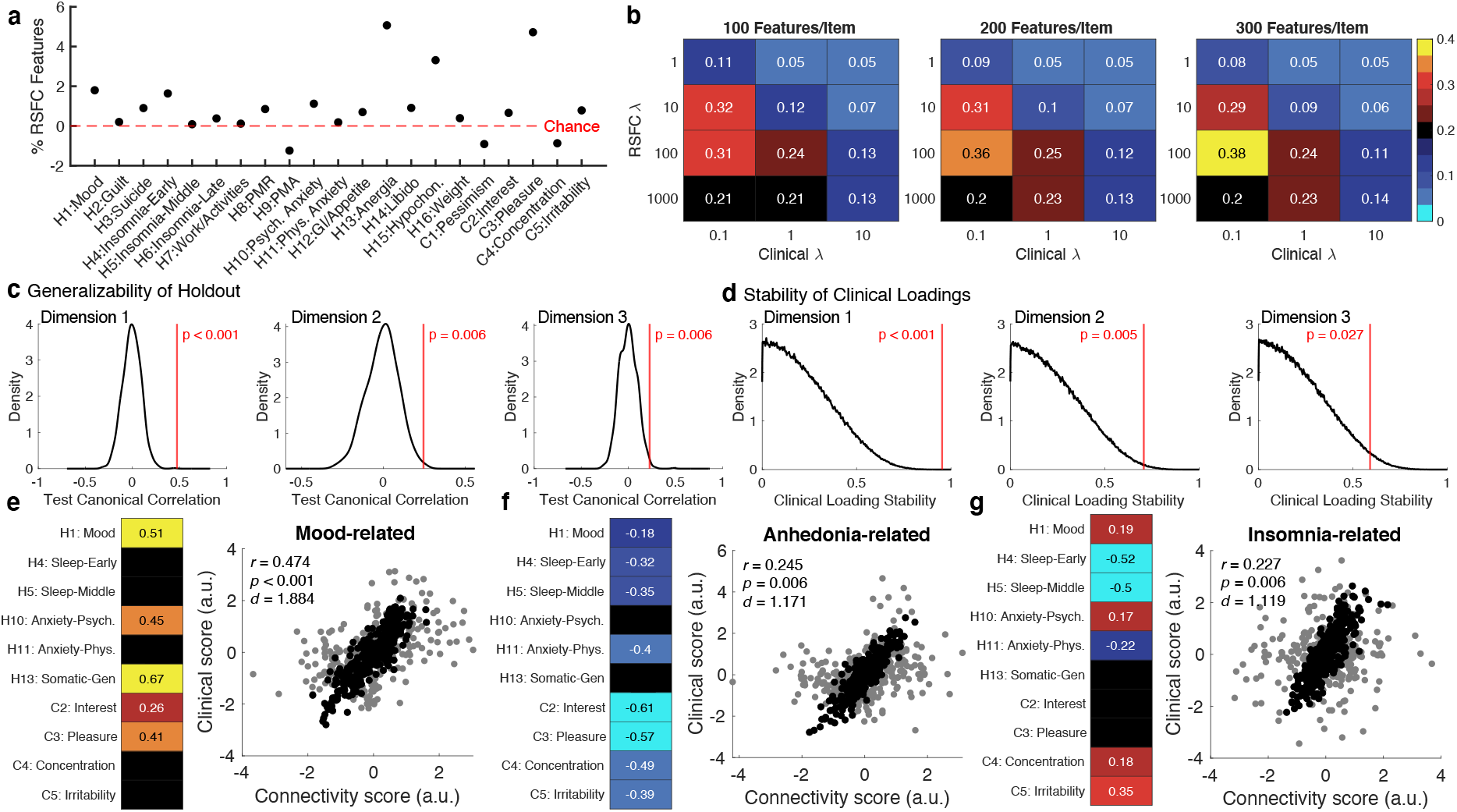
Three brain-behavior dimensions explaining individual differences in MDD. (**A**) On average, most clinical items have numerous resting-state functional connectivity (RSFC) features that exceed chance. Violin plots visualizing the number of RSFC features correlated with each clinical item at *p* < 0.05 using bootstrapping (1,000 replicates). Points indicate percentage of all 33,153 RSFC features that exceeded chance at *p* < 0.05, whole brain. The red bar indicates the chance number of clinical loadings, which has been standardized such that chance = 0. The first 16 items represent the Hamilton Rating Scale for Depression items, while the last 5 represent composite score from homologous items of the Beck Depression Inventory II and Inventory of Depressive Symptomatology. (**B**) Grid search hyperparameter optimization suggests that the optimal hyperparameters are: 300 RSFC features per clinical item, RSFC regularization λ (y-axes) = 100, and clinical regularization λ (x-axes) = 0.1. Heatmaps represent the average canonical correlation for canonical variate 1 when projected to data held-out during training within the inner fold (2/3 train, 1/3 test; 2,000 replicates). (**C**) The first three canonical variates significantly replicate in held-out data. Black kernel density estimations represent the histogram of permuted data (1,000 replicates), whereas the red line indicates the average test canonical correlation across 1,000 unshuffled replicates. For replicability in test data, the x-axis represents the canonical correlation of each canonical variate (CV) in test data at the optimal combination of hyperparameters identified in (B); greater canonical correlations in unshuffled data indicates a better fit of the regularized CCA model to test data. Corrected *p*-values (in red) were obtained using the null distribution. (**D**) The first three canonical variates have significantly stable clinical loadings. The x-axes represent the absolute correlation of clinical loadings; a higher score indicates greater agreement in clinical loadings across all training replicates. Black kernel density estimations represent the histogram of permuted training data (1,000 replicates), whereas the red line indicates the average training clinical loading correlation across 1,000 unshuffled replicates for each CV. Corrected *p*-values (in red) were obtained using the null distribution. (**E-G**) Results of the regularized CCA model, using the optimal combination of hyperparameters identified in Figure 1B, and using the entire sample (n = 328). Each figure section represents the following two images for each canonical variate (CV), from left to right: (I) A scatterplot visualizing the canonical correlation, with rho values displayed at the top. (II) significant clinical loadings, Bonferroni-corrected for the number of clinical items (*q* = 0.002). Black cells indicate that the clinical item was not statistically significant after correcting for multiple comparisons and heatmap values indicate the Pearson correlation coefficient of the clinical item severity and the CV scores in the clinical dimension. Higher values indicate a greater positive correlation between symptom severity and canonical variate score.

**Table 1:** Demographic information for all MDD participants analyzed during regularized CCA. BDI-II: 21-item Beck Depression Inventory II(49); IDS: 28-item Inventory of Depressive Symptomatology(50); HRSD: 17-item Hamilton Rating Scale for Depression(44); MDD: Major Depressive Disorder; SD: Standard Deviation.

|  | MDD |
| --- | --- |
| *n* | 328 |
| Female (%) | 215 (65.6%) |
| Age (SD) | 40.354 (12.047) |
| *n* HRSD (%) | 328 |
| HRSD (SD) | 22.119 (4.784) |
| *n* BDI-II (%) | 215 (65.5%) |
| BDI-II (SD) | 37.58 (11.76) |
| *n* IDS (%) | 113 (34.5%) |
| IDS (SD) | 38.248 (8.486) |

Next, we used regularized canonical correlation analysis to identify brain-behavior dimensions explaining individual differences in depressive symptoms, with cross-validation and testing in held-out data to reduce overfitting (for details, see Supplementary Methods, and for a schematic of this procedure, see **Supplementary Figure 2A)**. To this end, we first identified the optimal regularization parameters that maximized the canonical correlation of the first brain-behavior dimension (or canonical variate, CV1) in held-out data. The optimal combination involved 300 RSFC features and regularization parameters depicted in **Figure 1B**. These parameters were selected in 98/100 grid search replicates (outer training canonical correlation = 0.91 ± 0.01SD; outer test canonical correlation = 0.49 ± 0.06SD).

To test the statistical significance of CCA performance in unseen data (**Supplementary Figure S2B**), we generated a null distribution for canonical correlations in the test set by projecting coefficients modelled using intact training data and shuffled test replicates. The first three brain-behavior dimensions were statistically significant (**Figure 1C**), with canonical correlations in held-out test data ranging from r = 0.23 to 0.47, indicating large effect sizes compared (Cohen’s d = 1.12–1.88, p < 0.006). The fourth brain-behavior dimension in this model was not statistically significant in held-out test data (r = 0.08, p = 0.21).

To further validate these results, we also tested the stability of clinical symptom loadings generated during training for each brain-behavior dimension (1,000 replicates, **Supplementary Figure S2C**). To gauge the similarity of clinical symptom loadings across all training replicates, we calculated the absolute correlation between clinical symptom loadings for each dimension, such that higher absolute correlations indicated greater stability in clinical loadings across training replicates. Overall, the clinical symptom loadings were highly stable across training set replicates for the first three brain-behavior dimensions (**Figure 1D**; r = 0.59–0.95, p < 0.027 compared to shuffled data). In contrast, the clinical symptom loadings for the fourth brain-behavior dimension were relatively unstable (r = 0.36, p = 0.22 compared to shuffled data). Therefore, all subsequent analyses focused on the first three dimensions identified by regularized CCA.

Having identified three robust brain-behavior dimensions that were stable across training set replicates and reproducible in held-out data, we set out to characterize the clinical symptom domains captured in each dimension by examining the clinical symptom loadings for each dimension (i.e., Pearson’s correlations between each clinical symptom item and CV scores). The first brain-behavior dimension (CV1)—“mood-related connectivity”—explained individual differences in mood symptoms, anxiety, fatigue, and somatic symptoms, such that higher dimension scores were associated with greater symptom severity (**Figure 1E**). The second brain-behavior dimension (CV2)—“anhedonia-related connectivity”—explained individual differences in hedonic function (**Figure 1F)**, with no association with anxiety or fatigue and only a modest association with mood symptoms. The third brain-behavior dimension (CV3)—“insomnia-related connectivity”—explained individual differences in insomnia, especially difficulties with initiating and sustaining sleep (**Figure 1G**), as well as a moderate association with irritability and no association with anhedonia. Taken together, this analysis reveals three independent brain-behavior dimensions explaining individual differences in distinct symptom domains in depression.

To characterize the neurobiological basis of these dimensions, we extended the analysis of clinical symptom loadings above to examine RSFC feature loadings (whole brain, FDR corrected). To visualize within- and between-network trends, we ranked the top 25 brain regions for each dimension (CV) by summed *R*^2^ (**Figure 2A-C**). For the mood-related dimension 1, higher scores—indicating higher levels of depressed mood, anxiety, and fatigue—were associated with higher thalamic and basal ganglia RSFC and greater connectivity within the default mode network (**Figure 2A, Figure S3**). Higher mood-related dimension scores were also correlated with lower RSFC between the default mode network, cingulo-opercular network (especially the anterior insula), and frontoparietal control network (including the dorsolateral prefrontal cortex). For the anhedonia-related dimension 2, higher scores were associated with reduced levels of anhedonia; more severe symptoms were associated with higher RSFC within and between the cingulo-opercular control network and the dorsal and ventral attentional networks, as well as visual network areas, among others (**Figure 2B, Figure S4**). Lastly, for the insomnia-related dimension 3, higher scores— indicating milder difficulties with initiating and sustaining sleep and less irritability—were predominately associated with sensorimotor RSFC and RSFC between basal ganglia and the ventromedial prefrontal cortex (**Figure 2C, Figure S5**).

**Figure 2:**
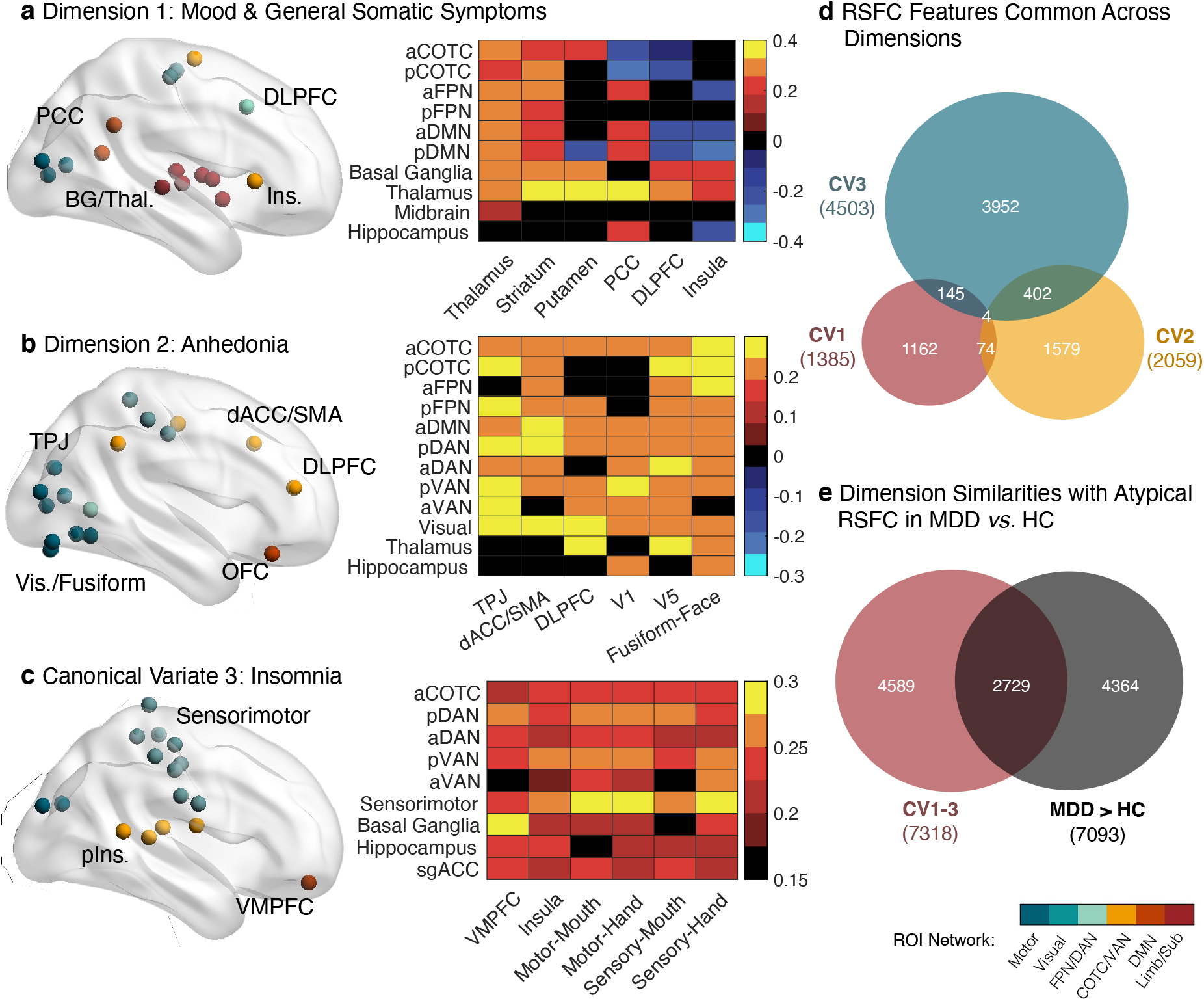
Resting-state functional connectivity correlates of three brain-behavior dimensions. **(A-C)** Results of the regularized CCA model, using the optimal combination of hyperparameters identified in Figure 1B, and using the entire sample (n = 328). Each figure section represents the following two images for each canonical variate (CV), from left to right: (I) The top 25 regions of interest associated with each canonical variate, identified as the sum of coefficients of determination for surviving features at FDR-*p* < 0.05, whole-brain. Only the right hemisphere is visualized, and regions of interest are color-coded by network. (II) Network-binned RSFC loadings for the top 25 regions of interest. Top regions of interest from the same network and region were binned to identify the mean RSFC loading (Pearson correlation coefficient, FDR-corrected) for intrinsic brain networks. Black cells indicate that no RSFC features remained after correcting for multiple comparisons. Higher values indicate a greater positive correlation between RSFC and canonical variate score. Canonical variate 1 was associated with depressed mood and general somatic symptoms, increased default mode/subcortical RSFC, and decreased default mode-cingulo-opercular RSFC. Canonical variate 2 was associated with anhedonia, and decreased cingulo-opercular, attentional, and visual RSFC. Canonical variate 3 was associated with insomnia and increased sensorimotor RSFC. (**D**) Venn diagram depicting the degree of overlap between the three canonical variates’ significant RSFC features. Only 4 RSFC features were significantly associated with individual symptom differences for all three canonical variates. (**E**) Venn diagram depicting the degree of overlap between the three canonical variates’ significant RSFC features and atypical RSFC relative to nondepressed controls. Abbreviations: aCOTC: prefrontal/insular nodes of the cingulo-opercular network; aDAN: prefrontal nodes of the dorsal attention network; aDMN: prefrontal nodes of the default mode network; aFPN: prefrontal nodes of the fronto-parietal network; aVAN: prefrontal nodes of the ventral attention network; BG: basal ganglia; C: composite clinical item; CV: canonical variate; dACC/SMA: dorsal anterior cingulate cortex and supplementary motor area; DLPFC: dorsolateral prefrontal cortex; H: Hamilton clinical item; HC: healthy controls; Ins.: Insula; MDD: major depressive disorder; OFC: orbitofrontal cortex; pCOTC: parieto-temporal nodes of the cingulo-opercular network; PCC: posterior cingulate cortex; pDAN: parieto-temporal nodes of the dorsal attention network; pDMN: parieto-temporal nodes of the default mode network; pFPN: parieto-temporal nodes of the fronto-parietal network; Phys.: physiological; pIns: posterior insula; Psych.: psychological; pVAN: parieto-temporal nodes of the ventral attention network; ROI: region of interest; RSFC: resting-state functional connectivity; Thal.: thalamus; TPJ: temporo-parietal junction; V1: primary visual cortex; V5: middle temporal visual area; Vis.: visual; VMPFC: ventromedial prefrontal cortex.

In our previous work using a similar approach to model heterogeneity in autism spectrum disorder, we found that distinct sets of RSFC features explained individual differences in three autism spectrum behavioral domains, but unexpectedly, most of these RSFC features fell within a normal range (37). Thus, we performed a similar analysis here, testing whether distinct RSFC features explained individual differences in each brain-behavior dimension and whether these differences were explained predominantly by RSFC features that were abnormal compared to healthy controls or by RSFC variation that fell within the normal range. This analysis showed that the vast majority of significant RSFC loadings were specific to one brain-behavior dimension (**Figure 2D**): 6,693 RSFC features were significantly correlated with just one brain-behavior dimension score (FDR corrected), but only 625 (8.5%) were significantly correlated with two or more brain-behavior dimensions. Next, we evaluated the extent to which the 7,318 RSFC features that were significantly correlated with at least one brain-behavior dimension score were also abnormal in MDD patients compared to a reference healthy control dataset comprising 461 individuals (266 females) with no history of depression. 37.3% of these RSFC features were abnormal in MDD patients compared to healthy controls (FDR corrected p < 0.05), but a majority (62.7%) fell within the normal range of variation in individuals without depression (**Figure 2E**). Together, these results confirm what was previously observed in autism spectrum disorder (37): individual differences in depressive symptoms are explained predominantly by variation in specific RSFC features that falls within the normal range and is associated with MDD symptoms only when it co-occurs with a distinct, largely non-overlapping set of abnormal RSFC features.

The results above define three independent brain-behavior dimensions that explain individual differences in mood symptoms, hedonic function, and insomnia in MDD patients that are stable across training replicates and reproducible in held-out data. To better understand the conditions necessary to generate this model, we tested how CCA model performance (first canonical correlation in held-out data) varies as a function of the sample size used in training and the effect size for correlations between individual RSFC features and individual correlations. We repeated the same hyperparameter tuning step identified in **Supplemental Figure 2A**, manipulating the sample size or the strength of univariate correlations between RSFC and clinical data included in the RCCA model (supplemental methods). We evaluated the stability of the optimal hyperparamter solution identified in the inner training/test fold, and the strength of the first canonical correlation in unseen data in the outer fold. We found that both sample size and the strength of univariate correlations influenced both the stability and replicability of the optimal hyperparameter solution as summarized in the Supplemental Results (**Table S1**). As expected, increasing the sample size and the strength of univariate correlations improved both the stability and performance of this modelling approach.

### Hierarchical Clustering Reveals Four Robust and Stable MDD Subtypes

Having identified three brain-behavior dimensions explaining individual differences in MDD symptoms, we used hierarchical clustering to identify relatively homogeneous subgroups of MDD patients in this hybrid brain-behavior space (**Figure 3A-B**). As in our previous work, we began by clustering MDD patients on their component scores for the first two brain-behavior dimensions (canonical variates) with the strongest correlations in Figure 1E-F. To identify the optimal number of clusters, we compared the Calinski-Harabasz (CH) criterion value—a measure of cluster quality based on the ratio of between-cluster variance to within-cluster variance—for clustering solutions involving 2–10 clusters (**Figure 3C**). To evaluate whether the degree of clustering in each solution exceeded that expected by chance, we generated a null distribution of CH criterion values in shuffled data over 1,000 replicates (**Figure 3D**; see Methods for details). This analysis revealed an optimal clustering solution involving four distinct subgroups of MDD patients (**Figure 3C**) defined by distinct patterns of mood- (CV1) and anhedonia-related functional connectivity (CV2). As in our previous work in MDD and autism spectrum disorder (2,37), individual MDD patients were not distributed into markedly discrete subgroups, but the degree of clustering observed in this solution was significantly greater than expected by chance (p = 0.004, **Figure 3D**). To further validate these subgroups, we assessed the stability of cluster assignments based on hierarchical clustering in bootstrapped subsamples of the data (2/3 of the full dataset across 1,000 replicates) compared to cluster assignments derived from hierarchical clustering on the full dataset. Overall, cluster assignments were stable across subsamples (**Figure 3E**), and the degree of cluster membership concordance for the four-cluster solution was significantly greater than expected by chance (p < 0.001, **Figure 3F**). Of note, we also tested for higher-order cluster solutions. Hierarchical clustering on all three dimensions did not reveal significant evidence of clustering in this space (**Supplementary Figure S6)**. This does not rule out the possibility of additional clusters, but in the dataset available to us, a four-cluster solution in a 2D space defined by the first two brain-behavior dimensions was optimal. Therefore, all subsequent analyses focused on this solution.

**Figure 3:**
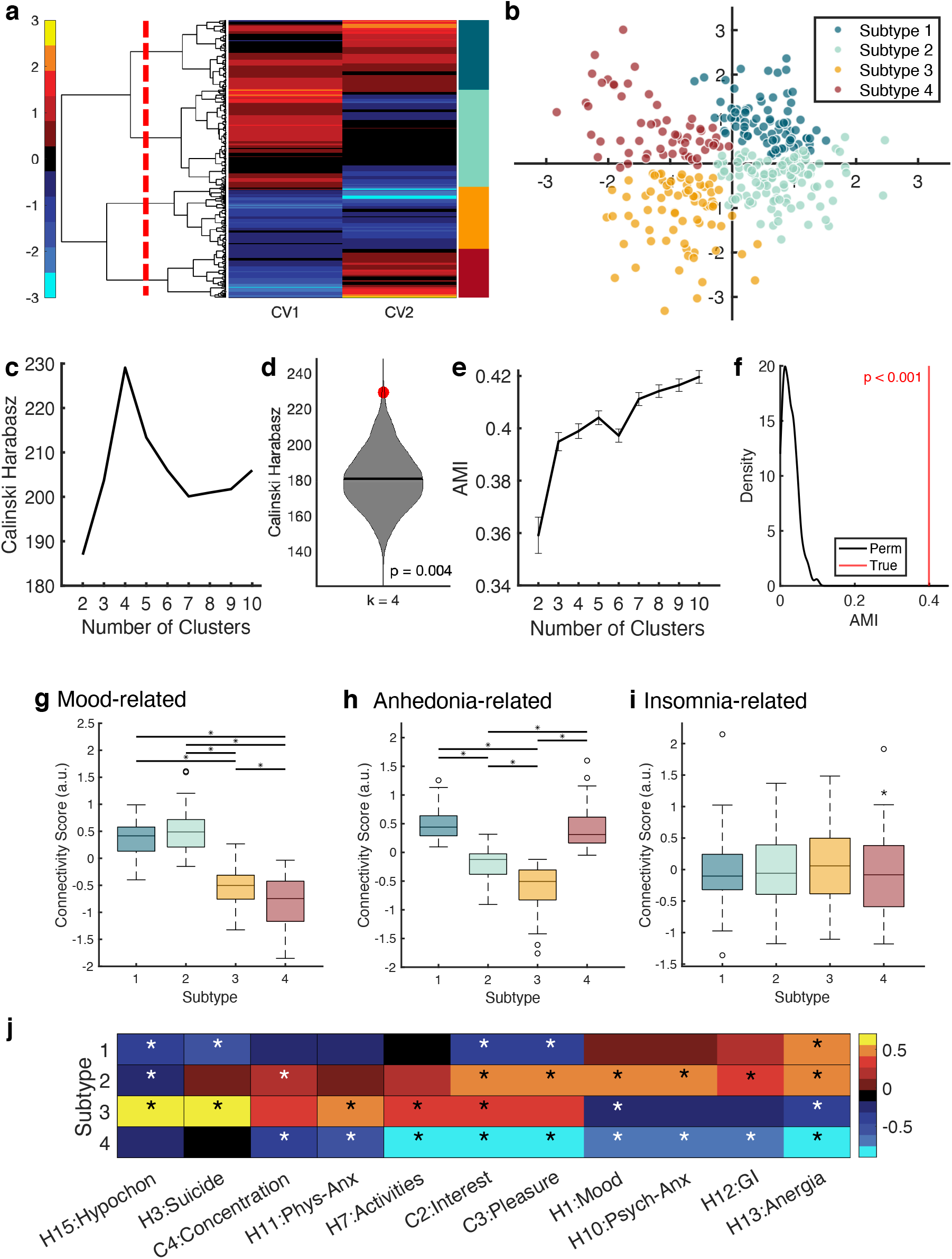
Hierarchical Clustering Reveals Four Robust and Stable MDD Subtypes. (**A**) Dendrogram visualizing the optimal *k* = 4 solution using the first two canonical variates. Heatmap values indicate participants’ canonical variate scores in the RSFC dimension (arbitrary units). (**B**) Scatterplot visualizing the four clusters by canonical variate score. The x- and y-axes represent the canonical variate scores 1 and two, respectively. (**C**) Calinski-Harabasz criteria values evaluating 2-10 cluster solutions when suing the first two canonical variates. Higher values indicate better performance in terms of the within- and between-cluster variance. The red asterisk indicates candidate hierarchical clustering solutions, which represents a peak in the Calinski-Harabasz criterion at *k* = 4. (**D**) Permutation testing indicates that the *k* = 4 clustering solution using the first two variates performs significantly better than that expected by chance. The violin plot represents the distribution of Calinski-Harabasz Criterion values for permuted training data; the red point indicates the criterion value for the unshuffled cluster solution. (**E**) Adjusted mutual information assessing the concordance of cluster membership in two-thirds subsamples of data relative to membership obtained using the entire sample (error bars = standard error of the mean). The elbow is at *k* = 3 clusters using the first two canonical variates. (**F**) Permutation testing the candidate optimal solution, *k* = 4 clusters, using the first two canonical variates. Cluster membership in two-thirds subsamples relative to the membership obtained using the complete sample for this potential clustering solution performed significantly greater than chance. The kernel density estimation represents the distribution of adjusted mutual information values in shuffled data, while the red line indicates the mean adjusted mutual information of the true sample. (**E-G**) Boxplots visualizing dimension scores for canonical variates 1 (E), 2 (F), and 3 (G). Asterisks indicate significant differences at FDR-*p* < 0.05. There were robust differences by subtype for the first two canonical variates. (**H**) Mean clinical severity by subtype for the Hamilton Rating Scale for Depression and composite scores; error bars represent the standard error of the mean. Asterisks represent subtype-specific significant differences (FDR-*p* < 0.05) relative to the severity of the entire sample. Asterisks indicate significant differences at FDR-*p* < 0.05. Abbreviations: C: composite score; CV: canonical variate; H: Hamilton Rating Scale for Depression item.

Next, we tested whether the four putative MDD subtypes identified above differed with respect to their clinical symptom profiles. The subtypes did not differ significantly with respect to age (Kruskal-Wallis χ^2^=1.026, *df*=3, *p*=0.795, ε²=0.005) or sex (χ^2^=4.502, *df*=3, *p*=0.212). However, multiple MDD symptoms varied by subtype (**Figure 3G-I, Supplementary Tables 2–3**), with mood, anxiety, somatic, and anhedonia symptoms differing most strongly. Subtypes 2 and 3 exhibited the most severe anhedonia symptoms and were also associated with elevated levels of suicidal ideation and anxiety compared to Subtypes 1 and 4. Subtype 1 was associated with low levels of anhedonia and high levels of anxiety and somatic symptoms such as appetite loss and fatigue. Subtype 4 was associated with less severe symptoms across a range of domains. As expected, these differences in clinical symptom profiles were associated with subtype-specific connectivity patterns for the mood- and anhedonia-related brain-behavior dimensions but not for the insomnia-related dimension. Specifically, Subtypes 1 and 2 were associated with relatively elevated mood-related connectivity scores (CV1)—predicting higher levels of mood and anxiety symptoms—while Subtypes 1 and 4 were associated with relatively elevated anhedonia-related connectivity scores (CV2)—predicting *lower* levels of anhedonia.

### Subtyping reveals distinct RSFC abnormalities and differences in TMS outcomes

Motivated by these findings, we concluded our analyses by evaluating the extent to which the four subtypes defined above were associated with distinct patterns of abnormal RSFC compared to healthy controls and with differences in treatment outcomes. As above, we identified RSFC features that were significantly abnormal in each subtype compared to a reference healthy control dataset comprising 461 individuals (266 females) with no history of depression (by Wilcoxon rank sum test, FDR corrected). To visualize the neuroanatomical distribution of abnormal connectivity in each subtype, we identified the 25 ROIs with the most subtype-specific connectivity profiles, plotted in **Figure 4A**. These nodes included the bilateral DLPFC, dorsal and ventral higher order visual areas, dorsal striatum, thalamus, and right orbitofrontal cortex. The most anhedonic Subtypes 2 and 3 were associated with hyperconnectivity in DLPFC-posterior FPN and DMN-thalamic networks, while subtypes with severe mood and anergia symptoms were associated with hypoconnectivity within and between most networks, except for thalamic hyperconnectivity (**Figure 4B-E**). Subtypes 1 and 4 were associated with the largest number of atypical RSFC features, with 9,214 and 5,692 significantly abnormal RSFC features, respectively, while subtypes 2 and 3 had fewer atypical features, with 3,790 and 2,962 significantly abnormal RSFC features, respectively. Subtypes with similar dimension scores had more shared atypical RSFC features: for example, subtypes 1 and 2 (associated with greater dimension 1 scores) shared 1,117 abnormal RSFC features, while subtypes 1 and 4 (associated with greater dimension 2 scores) shared 1,552 abnormal RSFC features. In summary, the four subtypes exhibited both common and distinct patterns of atypical RSFC, with the greatest divergence localized to striatal and thalamic regions.

**Figure 4:**
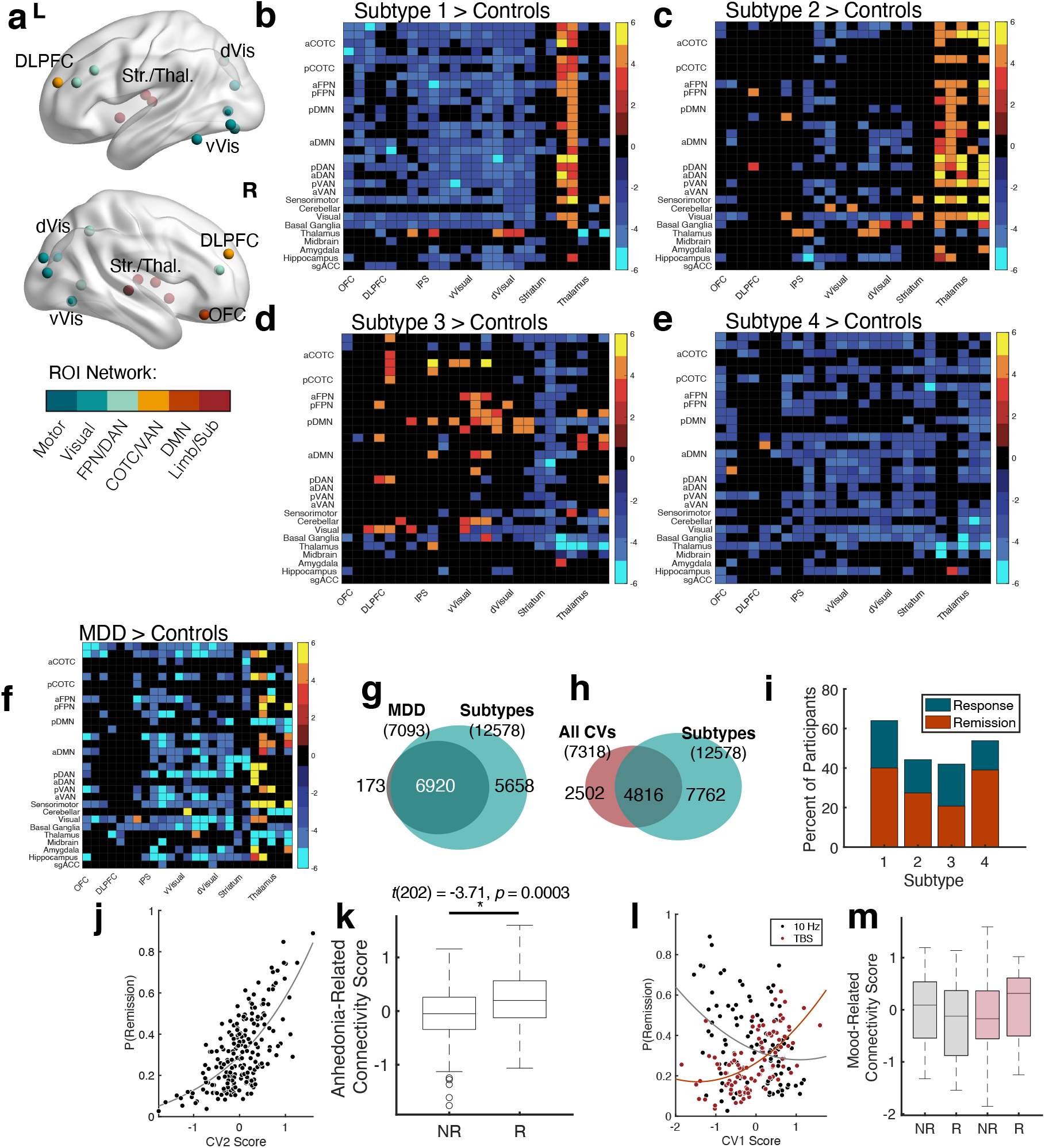
Subtyping reveals distinct RSFC abnormalities and differences in TMS outcomes. (**A**) The top 25 most distinct regions of interest in terms of subtype-specific atypical RSFC, identified using a Kruskal-Wallis test. (**B-E**) Network-binned RSFC loadings for the top 25 most divergent regions of interest across all four subtypes. ROIs from the same network and region were binned to identify the mean RSFC loading (*z*-score from a Wilcoxon rank sum test between subtypes and controls, FDR-corrected) for intrinsic brain networks. Black cells indicate that no RSFC features remained after correcting for multiple comparisons. Higher values indicate subtype hyperconnectivity relative to nondepressed controls. (**F**) Network-binned RSFC loadings representing atypical RSFC in all MDD participants relative to healthy controls, using the top 25 ROIs visualized in (B). (**G**) Venn diagram depicting the degree of overlap between significantly abnormal RSFC features present in the MDD *versus* healthy controls contrast, and RSFC features that were atypical in one or more subtypes. (**H**) Venn diagram depicted the degree of overlap between RSFC features that were significantly atypical in one or more subtypes and significantly associated with symptom dimensions (canonical variate loadings). (**I**) Differences in antidepressant response and remission to active rTMS (dorsolateral or dorsomedial rTMS, both 10 Hz and intermittent theta burst). Response was defined as ≥50% improvement from baseline to end of treatment on the Hamilton Rating Scale for Depression, and remission was defined as <9 on the same scale at the end of treatment. (**J-M**) Significant main effects and interactions from a binomial logistic regression predicting remission status. There was a significant main effect of canonical variate 2 score (**J-K**), such that a higher dimension score (lower anhedonia severity) resulted in a greater probability of remission, irrespective of stimulation site or treatment parameters. Furthermore, there was a significant interaction effect with canonical variate 1 (mood/somatic symptoms; **L-M**), such that dimension scores were differentially associated with 10 Hz and TBS remission. Abbreviations: aCOTC: anterior nodes of the cingulo-opercular task-control network; anterior nodes of the dorsal attention network; aDMN: anterior nodes of the default mode network; aFPN: anterior nodes of the fronto-parietal network; aVAN: anterior nodes of the ventral attention network; CV: canonical variate; DLPFC: dorsolateral prefrontal cortex; dVis: higher order visual, dorsal stream; IPS: intraparietal sulcus; NR: Non-Remitter; OFC: orbitofrontal cortex; Parahipp.: parahippocampal gyrus; pCOTC: posterior nodes of the cingulo-opercular task-control network; pDAN: posterior nodes of the dorsal attention network; pDMN: posterior nodes of the default mode network; pFPN: posterior nodes of the fronto-parietal network; pVAN: posterior nodes of the ventral attention network; R: Remitter; rTMS: repetitive transcranial magnetic stimulation; sgACC: subgenual cingulate cortexStr.: Striatum; TBS: theta burst stimulation; Thal.: thalamus; vVis: higher order visual, ventral stream.

Next, we sought to assess whether testing for abnormal functional connectivity within a given subtype revealed functional connectivity abnormalities that would not have been detected in a comparison of all MDD patients vs. healthy controls. In other words, do subtype-specific analyses provide more information about atypical RSFC? In a contrast of all MDD patients vs. healthy controls, we identified a range of abnormalities spanning the cingulo-opercular, frontoparietal, default mode and attention networks (**Figure 4F**). Only 2.4% of features significant in the MDD > control contrast were absent from subtype-specific contrasts (**Figure 4G**). Conversely, 45% of the features significant in one or more subtypes were absent from the MDD vs. control contrast, and nearly all (90.5%) these abnormal features were specific to only one subtype. These results indicate that subtype-specific contrasts captured atypical RSFC features associated with depression that would not have been identified without subtyping. Lastly, we tested whether subtype-specific atypical RSFC features overlapped with RSFC features with significant CV loadings, as we showed in **Figure 2E** that significant brain-behavior associations were not necessarily those identified in the MDD > control contrast. While there was some overlap (**Figure 4H**), many of the RSFC abnormalities did not significantly explain individual differences in symptoms (i.e. 39.6% of abnormal RSFCs were also significantly correlated with CV scores) or *vice versa* (61.7%), indicating that individual differences in depressive symptoms are still explained predominantly by variation in specific RSFC features that falls within the normal range, even after considering subtype-specific abnormalities.

Finally, we tested whether CVs or subtypes predicted responsivity to rTMS treatment. MDD participants in our sample received rTMS treatment in one of three studies(2,47,48). As each of these studies used different stimulatory parameters, we restricted the analysis to participants who received 4-6 weeks of once daily rTMS using intermittent theta burst (TBS) or 10 Hz stimulation over either the DLPFC or DMPFC (*n* treatment completers: DLPFC-10Hz=60, DLPFC-TBS=50, DMPFC-10Hz=49, DMPFC-TBS=45). Antidepressant response and remission to rTMS—irrespective of stimulation site or parameters—was significantly greater in the two subtypes associated with lower anhedonia (Response: χ^2^=4.671, *p*=0.031 Remission: χ^2^=5.568, *p*=0.018; **Figure 4I**). We investigated this further using logistic regression predicting remission status, with age and CV1-2 score as covariates; sex, stimulation site (DLPFC/DMPFC) and stimulatory parameter (10 Hz/TBS) as factors, and two-way interactions between CV scores, and treatment conditions (site/stimulatory parameter). Significant main effects for this model (χ^2^=29.722, *df*=10, *p*=0.001, **Table S4**) included the anhedonia dimension CV2 (estimate=1.417±0.570 SE, Odds Ratio(OR)=4.125 [1.350 12.607 95% CI], *z*=2.486, *p*=0.013, **Figure 4J-K**), and sex, such that females were more likely to remit than males (estimate=0.943±0.378 SE, OR=2.569 [1.350 12.607 95% CI], *z*=2.497, *p*=0.013). There was also a significant CV1*Stimulation Parameter interaction, such that 10Hz remitters had lower CV1 scores (less severe mood/somatic symptoms) and *vice versa* for TBS remitters (estimate=1.040±0.509 SE, OR=2.828 [1.043 7.670 95% CI], *z*=2.042, *p*=0.041, **Figure 4L-M**). Although this analysis was not designed to predict differential response to these two targets, we also tested whether there were any differences by stimulation site. There were no significant main or interaction effects by stimulation site (*p*>0.652). Consistent with our previous finding (2), the results indicate that favorable rTMS outcomes are associated with less severe anhedonia.

## DISCUSSION

In this extension of our previous work, we sought to develop and evaluate both dimensional and categorical solutions to parsing heterogeneity in depression using updated statistical approaches in a large, deeply phenotyped, single-site sample. The performance and stability of both regularized CCA and subsequent hierarchical clustering were significant. We also aimed to further assess co-occurring depressive symptoms and RSFC. The final CCA model yielded three dimensions explaining individual differences in mood/somatic symptoms associated with thalamic/DMN RSFC; anhedonia, correlated with visual and cingulo-opercular RSFC; and insomnia, associated with sensorimotor and posterior insula RSFC. Dimension scores clustered participants into four homogeneous subtypes, each with distinct clinical symptom profiles, abnormal RSFC patterns, and rTMS responsivity.

We also evaluated situations in which our approach may not yield robust or stable dimensions. At sample sizes < 150, a nested grid search approach poorly generalized to unseen data and did not stably converge on an optimal combination of model parameters. Furthermore, even in a large sample, this approach did not perform well in data with weak correlations between RSFC features and clinical symptoms (Supplemental Table 1). Although we did not assess this directly due to technical constraints, we would predict that these problems would be more pronounced in scenarios involving both smaller samples and weaker RSFC-clinical symptom correlations, and reliable models may also be more challenging to build in multi-site samples. Other situations that we did not assess may pose challenges for this approach. For example, univariate RSFC-symptom correlations could be affected by item range or distribution, especially if the sample consists of predominantly subclinical populations, or if there are multiple sites or raters administering scales, potentially amplifying heterogeneity in the data and introducing uncontrolled sources of noise. fMRI data quality and batch effects could also negatively impact the strength of these correlations, and ideally scans should be of long duration and with very little motion from a single scanner. Multi-echo fMRI may also help to resolve issues of data collection duration and quality (68); however such data has yet to be used in such an analysis.

CCA yielded three significantly stable and generalizable dimensions that resembled well-documented MDD symptom-brain associations. For CV1, more severe symptoms correlated with increased within-network DMN RSFC and decreased between-network DMN RSFC with the cingulo-opercular network. Numerous studies reports that negative self-referential processing in MDD is associated with DMN RSFC hyperconnectivity and hypoconnectivity between the DMN and cingulo-opercular (salience) networks(16,55,56). DMN functional connectivity is also implicated in sad mood induction (57) and elevated peripheral inflammatory markers in MDD(58). This CV was also associated with elevated thalamocortical RSFC; this circuitry is implicated in depressive symptomatology in animal models (59) and melancholic depression (60). Also consistent with previous literature, CV2, associated with anhedonia, was correlated with attentional and cingulo-opercular RSFC (31,61–63). Lastly, CV3 was associated with lower sensorimotor and posterior insula RSFC; previous research implicates these regions in aberrant homeostatic regulation in nondepressed (64–66) and depressed participants(67) with insomnia.

The brain-behavior dimensions and subtypes that we identified in this report resemble those identified in our previous work (2). For example, our previous study identified *two* brain-behavior dimensions explaining individual differences in anhedonia and in anxiety/insomnia/ mood symptoms, respectively, with four subtypes clustered in this space. In the present work, our analysis in a larger single-site sample with deeper clinical phenotyping identified *three* brain-behavior dimensions explaining individual differences in anhedonia, mood/anxiety, and insomnia, respectively (i.e. the anxiety/insomnia/mood dimension in our previous work was separable into two dimensions in this report), and again, we identified an optimal four-cluster subtyping solution in this space. At the same time, the results of these two analyses are not identical. Given the differences in the analyses, such as the inclusion of a nested grid search, bootstrapped feature selection and regularization, we would not necessarily expect identical results, but the similarities enhance confidence in the qualitative findings.

Despite the updates in our methodological approach, we also identified similarities in subtype-specific atypical RSFC features compared to our previous analyses(2). For example, we found subcortical hyperconnectivity in anhedonic subtypes and orbitofrontal hypoconnectivity in subtypes with severe fatigue in both analyses (Figure 4b-d). As we previously reported(2), subtypes with lower anhedonic symptoms were most responsive to either DMPFC or DLPFC rTMS. Previous rTMS studies implicate baseline anhedonia in excitatory rTMS response over the DMPFC(51) or DLPFC(28,68,69). Future studies are needed to: (a) prospectively predict responsivity to rTMS based on subtype assignment, and (b) systematically test other antidepressant interventions with the ultimate goal of identifying optimal interventions for individuals within each subtype, especially when responsivity to rTMS is suboptimal.

We also note several caveats of the current analysis. First, this study was not intended to be a replication of our previous findings as we included some participants from the original analysis(2). Instead, it represents an analytic extension with nearly threefold more participants scanned at the same site. Nevertheless, the current results support future MDD subtyping endeavors and replication attempts using RSFC data. Second, the CV and subtypes could be sensitive to dataset idiosyncrasies like study design or clinical assessments; for example, both age (70) and sex (71) influence normative and depressed RSFC features. Participants included in the current study were referred for rTMS, meaning they were treatment-resistant, taking psychotropic medications, and predominately moderate-to-severely depressed. Although our original analysis indicated that subtype assignment did not differ by medication status (2), modest RSFC changes associated with psychotropic medication use have been previously reported (72–78). Furthermore, differences in task-related activity and RSFC by MDD stage— subclinical MDD, first episode, and recurrent—are also beginning to be characterized(79–82). Future research on systematically testing each of these clinical factors and the extent to which they affect the brain-behavior relationships or subtypes is warranted. Third, the results are likely influenced by our choice of clinical inputs. Future studies would benefit from deeper clinical phenotyping of depression severity and specific symptom domains, perhaps by quantifying domain-specific impairments, like the Pittsburgh Sleep Questionnaire for insomnia/hypersomnia (83), or the Dimensional Anhedonia Rating Scale for deficits in reward processing and anhedonia (84).

To conclude, this study aimed to evaluate the performance of methods to develop neurobiologically-based MDD subtypes. Model performance and stability were significantly superior to chance. Furthermore, a four-cluster solution was the optimal solution for parsing heterogeneity categorically in our sample, and subtype membership was associated with rTMS response. These results represent an important step forward in assessing data-driven subtyping methods and provide evidence supporting the premise that regularized CCA is an effective tool to identify stable and generalizable associations between RSFC and behavior.

## Disclosures

KD was supported by a CIHR Banting Postdoctoral Fellowship and is currently supported by a University of Toronto Department of Psychiatry Academic Scholars Award. KD is listed as an inventor on Cornell University patent applications on neuroimaging biomarkers for depression that are pending or in preparation. JD reports research grants from CIHR, the National Institute for Mental Health, Brain Canada, the Canadian Biomarker Integration Network in Depression, the Ontario Brain Institute, the Klarman Family Foundation, the Arrell Family Foundation, and the Edgestone Foundation; reports travel stipends from Lundbeck and ANT Neuro, reports in-kind equipment support for investigator-initiated trials from MagVenture, and is an advisor for BrainCheck, NeuroStim TMS and Salience Neuro Health. FVR receives research support from CIHR, Brain Canada, Michael Smith Foundation for Health Research, Vancouver Coastal Health Research Institute, and Weston Brain Institute for investigator-initiated research. Philanthropic support from Seedlings Foundation. In-kind equipment support for this investigator-initiated trial from MagVenture. He has received honoraria for participation in advisory board for Janssen. ZJD has received research and equipment in-kind support for an investigator-initiated study through Brainsway Inc and Magventure Inc. He is also on the scientific advisory board for Brainsway Inc. His work has been supported by the National Institutes of Mental Health (NIMH), the Canadian Institutes of Health Research (CIHR), Brain Canada and the Temerty Family and Grant Family Foundations. DMB receives grant support from the Canadian Institutes of Health Research, the US National Institutes of Mental Health, Brain Canada, and the Temerty Family through the Centre for Addiction and Mental Health Foundation and the Campbell Research Institute. DMB received research support and in-kind equipment support for an investigator-initiated study from Brainsway. DMB was the site principal investigator for three sponsor-initiated studies for Brainsway. DMB received in-kind equipment support from Magventure for three investigator-initiated studies. DMB received medication supplies for an investigator-initiated trial from Indivior. DMB has participated in an advisory board meeting for Janssen and advisory board meeting for Welcony Inc. CL is supported by grants from the National Institute of Mental Health, the National Institute on Drug Abuse, the Rita Allen Foundation, the Klingenstein-Simons Foundation, the Brain and Behavior Research Foundation, the Hope for Depression Research Foundation, the Pritzker Neuropsychiatric Disorders Research Consortium, and the Wellcome Leap Multichannel Psych Program. CL has served as a consultant to Compass Pathways PLC, Delix Therapeutics, and Brainify.AI, and is listed as an inventor on Cornell University patent applications on neuroimaging biomarkers for depression that are pending or in preparation. The authors report no other financial relationships or conflicts.

