## Supplementary Material for "Dimensional and Categorical Solutions to Parsing Depression Heterogeneity in a Large Single-Site Sample"

### Supplemental Methods

#### *Participants*

Regarding inclusion and exclusion criteria(1–3), MDD participants were between 18-65 years old and met DSM-5 diagnostic criteria(4) for a current major depressive episode. Participants with psychiatric comorbidities were included in the study, provided that the comorbid pathology was secondary to the major depressive episode, as assessed by a licensed study psychiatrist. MDD participants were excluded from the study if they had any psychotic disorder; active suicidal intent; substance abuse or dependence in the last three months; substantial neurological pathology; or had any contraindications to rTMS or MRI.

#### *Treatment Design*

All three studies involved rTMS treatment, although the stimulation site and/or stimulation parameters varied slightly. For participants included in Drysdale et al.(1), all received open-label rTMS (10 Hz or intermittent theta burst stimulation [iTBS]) over the bilateral dorsomedial prefrontal cortex (DMPFC), guided using structural MRI (see Bakker et al.(5) for more details). Participants were treated once daily for four weeks (20 rTMS sessions total). Antidepressant response was defined as ≥50% improvement from baseline to end of treatment on the 17-item Hamilton Rating Scale for Depression (HDRS), and remission was defined as an end of treatment score on the HDRS of ≤7. For participants from Blumberger et al.(2), participants were randomized to receive either once-daily 10 Hz left dorsolateral prefrontal cortex stimulation (LDLPFC) or once-daily iTBS over the LDLPFC. LDLPFC stimulation was performed under MRI guidance, and participants received 4-6 weeks of treatment (20-30 sessions). Antidepressant response was defined as ≥50% improvement from baseline to end of treatment on the 17-item Hamilton Rating Scale for Depression (HDRS), and remission was defined as an end of treatment score on the HDRS of ≤7. Lastly, for participants from Dunlop et al.(3), participants were enrolled in a double-blind, three-arm study of DMPFC-rTMS. Participants were randomized either: 20 Hz bilateral DMPFC-rTMS; 1 Hz bilateral DMPFC-rTMS; or placebo DMPFC-rTMS. All participants received twice-daily treatments over three weeks, for a total of thirty rTMS sessions. Antidepressant response was defined as ≥35% improvement from baseline to end of treatment on the 17-item Hamilton Rating Scale for Depression (HDRS), and remission was defined as an end of treatment score on the HDRS of ≤7.

*Neuroimaging Acquisition and Preprocessing*

All neuroimaging was acquired at Toronto Western Hospital (Toronto, Canada) on a 3 Tesla GE Signa HDx equipped with an eight-channel phased-array head coil. The neuroimaging acquisition parameters have been described in detail elsewhere(6). Briefly, all participants underwent at 30-minute protocol that involved a T1-weighted high-resolution anatomical scan (fast spoiled gradient-echo image; TE=12ms, TI=300ms, flip angle=20°, 116 sagittal slices, thickness=1.5mm, no gap, 256x256 matrix, FOV=240mm) and a 10-minute T2*-weighted resting-state functional MRI in the eyes closed condition (TE=30ms, TR=2000ms, flip angle=85°, 32 axial slices, thickness=5mm, no gap, 64x64 matrix, FOV=220mm, 300 volumes).

Resting-state scans were preprocessed using the Analysis of Functional Neuroimages (AFNI) software package(7). Preprocessing was identical to that described in Drysdale et al.(1) Briefly, all scans underwent standard procedures for slice-timing correction, spatial smoothing (6mm full-width half-maximum Gaussian kernel), and temporal bandpass filtering (0.01-0.1 Hz). Denoising steps included linear and quadratic detrending and removal of signals related to motion (six demeaned time series, its first temporal derivatives) and physiological/non-neuronal sources using ANATICOR(8) via the "AFNI_proc.py" command(9). We also censored volumes and their preceding/following volumes if its Euclidean Distance was greater than 0.3 mm. All scans were co-registered to the individual's high resolution T1-weighted scan, and then nonlinearly transformed to the Montreal Neurological Institute standardized space. This resulted in residual time series for each scan, preprocessed and denoised and in standardized space, with which to extract functional connectivity.

The functional parcellation by Power and colleagues(10) was used to extract whole-brain functional connectivity from preprocessed and denoised data. We used the same 258 regions of interest (ROI) described in Drysdale et al.(1) for all subsequent analyses (Supplementary Figure 1), resulting in 33,153 RSFC features. Resting-state functional connectivity was defined as the Fisher-*z* transformed Pearson Correlation Coefficient of an ROI pair's time series.

*Clinical Composite Scores*

To generate composite scores, we *z*-normalized each BDI-II and IDS clinical item. Clinical normalization was conducted using only data from the training set, using the mean and variance to standardize the test set. As the BDI-II and IDS were acquired for separate subsets of participants, we combined the appropriate items once they were standardized (i.e., mean=0, variance=1). In the case of concentration and decision-making for participants with BDI-II data, we calculated the mean of the two items after normalizing each individual item encompassing this symptom to generate a single score per participant.

*RSFC Feature Selection*

We performed bootstrapped feature selection using data partitioned to the training set to identify the most relevant RSFC features. Once the 21 clinical items (16 HRSD, 5 composite) were normalized, we partitioned and *z*-normalized RSFC data using the method described above. Similar to that described in Grosenick et al.(11), stabilized feature selection involved parametrically correlating each of the 33,153 RSFC features to each clinical item, using bootstrapping with replacement. Feature selection during hyperparameter optimization (Figure S2A) used 100 bootstrapping replicates, while feature selection to assess the generalizability and stability of CCA dimensions used 1,000 bootstrapping replicates. 21 whole brain rank lists, one for each clinical item, were generated using the absolute mean Pearson's correlation coefficient obtained during bootstrapping. Bootstrapped feature selection in this manner was performed for each training iteration, and for the final RCCA model using the entire dataset.

To determine whether RSFC and clinical associations using mass univariate statistics were robust, we calculated the number of RSFC correlations retained at four significance thresholds ($\alpha = 0.05$, 0.01, 0.005 and 0.001). Mean Pearson's Correlation Coefficients, obtained using the entire sample and 1,000 bootstrapping replicates, were transformed into *z*-values using the inverse cumulative distribution function of the normal distribution. The number of RSFC features of each clinical item retained for each $\alpha$ threshold were visualized both for correlations using the entire sample, and across all 1,000 training iterations.

*Hyperparameter Tuning*

Once RSFC rank lists were generated for a given inner fold iteration, we proceeded with training L2-norm regularized CCA using combinations of three hyperparameters: the number of RSFC features used during RCCA from each rank list (100, 200, 300 features per rank list, with duplicate features removed), and the RSFC and clinical regularization terms λ1 and λ2, respectively. λ1 was tested at 1, 10, 100 and 1000, while λ2 was tested at 0.1, 1, and 10. For each inner fold, this step generated 36 different combinations of hyperparameters for each iteration. Similar to Grosenick et al.(11), we selected the optimal combination of hyperparameters using the median canonical correlation in unseen data, which we then evaluated further in the outer fold test set.

*Orthogonal Deflation Procedure*

Sequential orthogonal deflation aims to minimize the impact of previously estimated canonical variates. After evaluating the first canonical variate and identifying the optimal combination of hyperparameters and number of RSFC feature, we performed orthogonal deflation to generate new whole-brain RSFC and clinical matrices for the training and test sets with which to assess canonical correlations and clinical loadings for the subsequent canonical variate. This process was repeated sequentially, provided that all previously estimated canonical variates significantly generalized to unseen data and had significantly stable clinical loadings.

After identifying and generating the canonical coefficients for the rank-*k* canonical variate via singular value decomposition, we defined the canonical components (scores) as:

$$U_k = X_k A_k \text{ and } W_k = Y_k B_k$$

Where $U_k$ and $W_k$ represent the rank-*k* canonical components for the RSFC and clinical matrices for the training data respectively; $A_k$ and $B_k$ are the canonical coefficients for canonical variate *k*; and $X_k$ and $Y_k$ are the rank-*k* approximation to the data matrices $X$ (RSFC; selected features only) and $Y$ (clinical). The $k + 1$ deflation is given by:

$$X_{k+1} = (I - P_{Uk})X_k = X_k - U_k\,(U_k^T\,U_k)^{-1}U_k^T\,X_k$$

$$Y_{k+1} = (I - P_{Wk})Y_k = Y_k - W_k\,(W_k^T\,W_k)^{-1}W_k^T\,Y_k$$

Where $Pu_k$ and $Pw_k$ represent the orthogonal projection matrices for the rank-$k$ canonical variate. Further, due to canonical coefficient shrinkage from L2-norm regularization(12), we also need to "un-shrink" each deflation estimate by estimating the scalar coefficients $\beta_{Uk}$ and $\beta_{Wk}$, obtained using univariate regression models: $X_k = \beta_{Uk}\, P_{Uk}\, X_k + \varepsilon_{Uk}$ and $Y_k = \beta_{Wk}\, P_{Wk}\, Y_k + \varepsilon_{Wk}$. As a result, the orthogonal deflation step is:

$$X_{k+1} = (I - P_{Uk})X_k = X_k - \beta_{Uk}U_k\,(U_k^T\,U_k\,)^{-1}U_k^T\,X_k$$
$$Y_{k+1} = (I - P_{Wk})Y_k = Y_k - \beta_{Wk}W_k\,(W_k^T\,W_k\,)^{-1}W_k^T\,Y_k$$

Lastly, to estimate the effect of canonical variate $k$ whole-brain (i.e., outside of selected RSFC features), we orthogonally deflated all 33,153 RSFC features using the projection matrix and scalar coefficient generated using RSFC features retained to estimate the coefficients for rank $k$ canonical variate:

$$Z_{k+1} = (I - P_{Uk})Z_k = Z_k - \beta_{Uk}U_k\,(U_k^T\,U_k\,)^{-1}U_k^T\,Z_k$$

Where $Z_k$ and $Z_{k+1}$ represent the whole-brain RSFC matrices for the rank-$k$ and rank-$k+1$ canonical variates. Orthogonal deflation of test data used the coefficients and projection matrices estimated using the training dataset.

*Assessing the effect of sample characteristics on hyperparameter training*

Lastly, we tested the impact of two sample characteristic variables on the stability and performance of hyperparameter tuning: sample size and the strength of RSFC-clinical correlations. To investigate this impact, we performed the methods outlined in Supplemental Figure 2A, except for the number of outer fold iterations to minimize computational considerations (10 iterations instead of 100). For sample size, we retained a random subset of participants with which to subdivide into the outer and inner folds, at 50 participant increments between n = 50 and n = 300. Next, for the strength of associations between clinical and RSFC data, we excluded RSFC features from training if they exceeded $\rho = 0.10$, 0.15, 0.20, or 0.25 on one or more correlations during bootstrapped feature selection. We evaluated two aspects of hyperparameter tuning to determine the performance of these scenarios: first, the stability of the optimally selected combination of hyperparameters across the ten outer fold iterations, and second, the performance of the first canonical variate in the held-out data in the outer fold.

### Supplementary Results

*Regularized CCA Yields Significant and Stable Symptom-Brain Dimensions*

Prior to regularized CCA optimization, we next examined the strength of correlations between clinical symptoms and RSFC. To ensure that the strength of associations between symptoms and RSFC data did not significantly differ between the 16 HRSD items and 5 composite scores, we compared the number of surviving RSFC features at four alpha thresholds (0.05, 0.01, 0.005 and 0.001) when we performed bootstrapped feature selection in 1,000 two-thirds subsample replicates. The strength of correlations between HRSD and composite scores did not significantly differ between the 16 HRSD items and five composite scores at all four thresholds (FDR-$p < 0.05$). The effect sizes of symptom-RSFC correlations were small to medium; none of the correlation coefficients exceeded |0.3| or a coefficient of determination of 0.07.

We conducted a grid search to identify the optimal combination of regularization parameters that maximized the canonical correlation of canonical variate (CV) 1 in unseen data. (100 replicates; **Figure S2A**). Three trends emerged when we optimized the regularization parameters using canonical variate 1 performance in test data (**Figure S3**). First, the optimal number of input features for all canonical variates assessed was the highest value we tested (300 features per rank list). That said, lowering the number of input features while keeping λ1 and λ2 stable had a small effect on the mean canonical correlation for CV1 extracted from test data. Second, regularization for RSFC data was generally higher than the clinical data, and there were clearly delineated local maxima at the λ1 values assessed while keeping the other two hyperparameters stable. Lastly, the optimal λ2 for all CVs assessed was among the lowest value tested ($\lambda2 = 0.1$); increasing this penalty on clinical data sharply decreased the generalizability of CVs, particularly when $\lambda2 > 1$.

*Sample characteristics impact the stability and performance of hyperparameter tuning*

We also manipulated three sample characteristics in order to better understand how these factors influence hyperparameter selection. We assessed the effect of sample size, strength of clinical-RSFC correlations, and restricted clinical item range during nested hyperparameter selection (**Figure S2A**). We measured the impact of these factors on the canonical correlation of test data in the outer fold, and how stable the optimal combination of hyperparameters were across 10 iterations (**Table S1**). Regarding sample size, both the stability of the optimal hyperparameter combination and the average test canonical correlation steadily increased as sample size increased. The strength of clinical-RSFC correlations impacted both the degree of regularization and the average test canonical correlation, such that as the strength of associations increased, the regularization and canonical correlation of the first dimension increased. We also note that, although this was not tested in this dataset, datasets acquired using multiple scanners is a sample characteristic that ought to be evaluated in future analyses as it likely impacts the stability and performance of such models. In sum, increased sample size and the strength of correlations increased the average test canonical correlation for the first dimension.

### Supplementary Tables

**Supplementary Table 1:** Impact of sample size, clinical item range, and strength of clinical-resting state functional connectivity (RSFC) correlations on CCA optimization across ten outer fold iterations. The number of RSFC features, RSFC and clinical regularization signify the most frequently selected hyperparameter solution during training and testing in the inner fold, and the proportion indicates how frequently this solution was identified across the ten iterations. The median and standard deviation report the canonical correlation for the first dimension in the outer test set using the optimal solution.

| | Number of RSFC features | RSFC Regularization | Clinical Regularization | Proportion | Median | Standard Deviation |
|---|---|---|---|---|---|---|
| *Original* | 300 | 100 | 0.1 | 0.98 | 0.487 | 0.057 |
| *Sample Size* | | | | | | |
| n = 50 | - | - | - | 0 | -0.075 | 0.163 |
| n = 100 | 300 | 1 | 10 | 0.2 | 0.088 | 0.188 |
| n = 150 | 300 | 100 | 0.1 | 0.4 | 0.263 | 0.117 |
| n = 200 | 300 | 100 | 0.1 | 0.5 | 0.331 | 0.135 |
| n = 250 | 300 | 100 | 0.1 | 0.7 | 0.372 | 0.097 |
| n = 300 | 300 | 100 | 0.1 | 1 | 0.491 | 0.072 |
| *Strength of Correlations* | | | | | | |
| $\rho < 0.10$ | - | - | - | 0 | 0.066 | 0.094 |
| $\rho < 0.15$ | 100 | 0.1 | 0.1 | 0.8 | 0.370 | 0.105 |
| $\rho < 0.20$ | 200 or 300 | 10 | 0.1 | 0.5 | 0.457 | 0.089 |
| $\rho < 0.25$ | 300 | 100 | 0.1 | 0.7 | 0.462 | 0.074 |

**Supplementary Table 2:** Post-hoc differences (Kruskal-Wallis) in baseline severity by subtype. A) Indicates the descriptive statistics (mean and standard deviations [SD]) for each subtype, and Kruskal-Wallis omnibus test, FDR-corrected. B) Indicates post-hoc tests (Wilcoxon rank-sum) for all significant tests in (A). p-values are FDR-corrected.

**A**

| Symptom | Subtype 1 | | Subtype 2 | | Subtype 3 | | Subtype 4 | | Z-Score | FDR-p |
|---|---|---|---|---|---|---|---|---|---|---|
| | Mean | SD | Mean | SD | Mean | SD | Mean | SD | | |
| Low Mood | 2.83 | 0.78 | 3.20 | 0.64 | 2.55 | 0.80 | 2.14 | 1.05 | 59.49 | 0.0000 |
| Guilt/Punishment | 2.07 | 0.90 | 2.25 | 0.92 | 2.32 | 0.81 | 1.71 | 0.99 | 16.48 | 0.0011 |
| Suicide | 0.83 | 0.87 | 1.35 | 1.05 | 1.99 | 1.1 | 1.29 | 1.12 | 41.43 | 0.0000 |
| Early Insomnia | 1.12 | 0.91 | 1.38 | 0.83 | 1.53 | 0.77 | 1.00 | 0.94 | 15.82 | 0.0014 |
| Mid Insomnia | 0.62 | 0.82 | 1.01 | 0.89 | 1.21 | 0.76 | 0.81 | 0.74 | 21.14 | 0.0001 |
| Late Insomnia | 0.40 | 0.74 | 0.79 | 0.87 | 0.88 | 0.80 | 0.52 | 0.73 | 19.91 | 0.0002 |
| Work/Activities | 3.13 | 0.58 | 3.31 | 0.64 | 3.45 | 0.69 | 2.55 | 0.99 | 41.97 | 0.0000 |
| PMR | 0.72 | 0.84 | 0.66 | 0.76 | 0.44 | 0.71 | 0.41 | 0.68 | 10.00 | 0.0195 |
| PMA | 0.42 | 0.57 | 0.30 | 0.55 | 0.40 | 0.66 | 0.29 | 0.59 | 4.04 | 0.2568 |
| Psych Anxiety | 2.22 | 0.82 | 2.51 | 0.84 | 1.88 | 0.91 | 1.45 | 0.99 | 52.73 | 0.0000 |
| Somatic Anxiety | 1.44 | 0.92 | 1.77 | 0.86 | 2.15 | 1.10 | 1.16 | 0.99 | 38.19 | 0.0000 |
| Appetite | 1.02 | 0.85 | 1.13 | 0.82 | 0.60 | 0.72 | 0.31 | 0.50 | 48.05 | 0.0000 |
| General Somatic | 1.74 | 0.47 | 1.74 | 0.48 | 1.26 | 0.47 | 0.98 | 0.55 | 95.47 | 0.0000 |
| Libido | 1.72 | 0.59 | 1.63 | 0.68 | 1.36 | 0.82 | 1.33 | 0.85 | 15.13 | 0.0019 |
| Hypochondriasis | 0.50 | 0.69 | 0.68 | 0.80 | 1.67 | 1.12 | 1.28 | 1.07 | 59.12 | 0.0000 |
| Weight | 0.44 | 0.79 | 0.38 | 0.72 | 0.21 | 0.53 | 0.02 | 0.13 | 17.43 | 0.0008 |
| Total HAMD Score | 21.23 | 2.86 | 24.10 | 3.70 | 23.88 | 4.69 | 17.24 | 5.23 | 82.46 | 0.0000 |
| Pessimism (Future) | -0.14 | 0.97 | 0.17 | 0.97 | 0.29 | 0.87 | -0.49 | 1.05 | 23.12 | 0.0001 |
| Interest/Involvement | -0.33 | 0.91 | 0.41 | 0.83 | 0.35 | 0.84 | -0.80 | 0.99 | 76.44 | 0.0000 |
| Pleasure/Enjoyment | -0.41 | 0.86 | 0.50 | 0.76 | 0.36 | 0.85 | -0.86 | 0.98 | 89.88 | 0.0000 |
| Decision-Making | -0.31 | 0.92 | 0.24 | 0.82 | 0.31 | 0.79 | -0.45 | 0.92 | 37.57 | 0.0000 |
| Irritability | -0.35 | 0.83 | 0.21 | 1.00 | 0.31 | 1.07 | -0.32 | 0.90 | 28.52 | 0.0000 |

**B**

| Symptom | Subtype 1>2 | | Subtype 1>3 | | Subtype 1>4 | | Subtype 2>3 | | Subtype 2>4 | | Subtype 3>4 | |
|---|---|---|---|---|---|---|---|---|---|---|---|---|
| | z | p | z | p | z | p | z | p | z | p | z | p |
| Low Mood | -3.44 | 0.00 | 2.41 | 0.03 | 4.09 | 0.00 | 5.53 | 0.00 | 6.73 | 0.00 | 2.05 | 0.06 |
| Guilt | -1.50 | 0.16 | -1.56 | 0.15 | 2.21 | 0.04 | -0.13 | 0.90 | 3.56 | 0.00 | 3.57 | 0.00 |
| Suicide | -3.41 | 0.00 | -6.25 | 0.00 | -2.40 | 0.03 | -3.92 | 0.00 | 0.34 | 0.77 | 3.46 | 0.00 |
| Early Insom. | -2.04 | 0.06 | -2.95 | 0.01 | 0.76 | 0.49 | -1.27 | 0.24 | 2.60 | 0.02 | 3.37 | 0.00 |
| Mid Insomnia | -3.03 | 0.01 | -4.38 | 0.00 | -1.74 | 0.11 | -1.45 | 0.18 | 1.38 | 0.20 | 2.91 | 0.01 |
| Late Insomnia | -3.32 | 0.00 | -4.06 | 0.00 | -1.27 | 0.24 | -0.84 | 0.44 | 1.90 | 0.08 | 2.67 | 0.01 |
| Work/Act. | -2.20 | 0.04 | -3.36 | 0.00 | 3.75 | 0.00 | -1.68 | 0.12 | 5.17 | 0.00 | 5.39 | 0.00 |
| PMR | 0.33 | 0.77 | 2.281 | 0.03 | 2.28 | 0.03 | 2.17 | 0.04 | 2.18 | 0.04 | 0.12 | 0.90 |
| Psych Anx. | -2.84 | 0.01 | 2.19 | 0.04 | 4.51 | 0.00 | 4.78 | 0.00 | 6.39 | 0.00 | 2.56 | 0.02 |
| Som. Anxiety | -2.31 | 0.03 | -4.59 | 0.00 | 1.63 | 0.13 | -3.22 | 0.00 | 3.67 | 0.00 | 5.24 | 0.00 |
| Appetite | -0.87 | 0.43 | 3.13 | 0.00 | 5.04 | 0.00 | 4.26 | 0.00 | 6.14 | 0.00 | 2.33 | 0.03 |
| Somatic | -0.17 | 0.89 | 5.72 | 0.00 | 7.17 | 0.00 | 6.33 | 0.00 | 7.81 | 0.00 | 2.92 | 0.01 |
| Libido | 0.93 | 0.40 | 3.05 | 0.01 | 3.02 | 0.01 | 2.41 | 0.03 | 2.41 | 0.03 | 0.16 | 0.89 |
| Hypochon. | -1.50 | 0.16 | -6.52 | 0.00 | -4.44 | 0.00 | -5.98 | 0.00 | -3.57 | 0.00 | 2.00 | 0.06 |
| Weight | 0.34 | 0.77 | 1.81 | 0.09 | 3.85 | 0.00 | 1.64 | 0.13 | 3.77 | 0.00 | 2.63 | 0.02 |
| Total HAMD | 7.77 | 0.00 | 5.29 | 0.00 | -6.74 | 0.00 | -0.97 | 0.90 | -10.99 | 0.00 | -9.15 | 0.00 |
| Pessimism | -2.36 | 0.03 | -3.06 | 0.01 | 1.85 | 0.09 | -0.94 | 0.39 | 3.55 | 0.00 | 4.22 | 0.00 |
| Interest | -5.85 | 0.00 | -5.28 | 0.00 | 2.80 | 0.01 | -0.77 | 0.48 | 6.89 | 0.00 | 6.19 | 0.00 |
| Pleasure | -6.43 | 0.00 | -4.00 | 0.00 | 3.09 | 0.00 | 2.67 | 0.01 | 8.04 | 0.00 | 6.30 | 0.00 |
| Concentration | -3.93 | 0.00 | -3.28 | 0.00 | 1.19 | 0.27 | 0.44 | 0.71 | 5.05 | 0.00 | 4.79 | 0.00 |
| Irritability | -3.87 | 0.00 | -3.54 | 0.00 | 1.15 | 0.29 | -0.38 | 0.75 | 3.94 | 0.00 | 3.57 | 0.00 |

**Supplementary Table 3:** FDR-Corrected p-values for Wilcoxon rank sum tests comparing subtype severity for each item relative to the full MDD sample.

| Symptom | Subtype 1 | Subtype 2 | Subtype 3 | Subtype 4 |
|---|---|---|---|---|
| Low Mood | 0.743 | 0.000 | 0.032 | 0.000 |
| Guilt/Punishment | 0.633 | 0.217 | 0.217 | 0.007 |
| Suicide | 0.001 | 0.990 | 0.000 | 0.727 |
| Early Insomnia | 0.194 | 0.372 | 0.052 | 0.056 |
| Mid Insomnia | 0.009 | 0.412 | 0.020 | 0.452 |
| Late Insomnia | 0.012 | 0.243 | 0.052 | 0.298 |
| Work/Activities | 0.405 | 0.168 | 0.007 | 0.000 |
| PMR | 0.231 | 0.316 | 0.168 | 0.167 |
| PMA | 0.256 | 0.508 | 0.718 | 0.388 |
| Psych Anxiety | 0.589 | 0.000 | 0.084 | 0.000 |
| Somatic Anxiety | 0.100 | 0.500 | 0.000 | 0.003 |
| Appetite | 0.122 | 0.004 | 0.053 | 0.000 |
| General Somatic | 0.003 | 0.000 | 0.001 | 0.000 |
| Libido | 0.080 | 0.316 | 0.102 | 0.100 |
| Hypochondriasis | 0.001 | 0.044 | 0.000 | 0.059 |
| Weight | 0.178 | 0.243 | 0.454 | 0.004 |
| Pessimism (Future) | 0.217 | 0.211 | 0.044 | 0.006 |
| Interest/Involvement | 0.004 | 0.002 | 0.003 | 0.000 |
| Pleasure/Enjoyment | 0.007 | 0.000 | 0.154 | 0.000 |
| Concentration/Decision-Making | 0.051 | 0.014 | 0.081 | 0.001 |
| Irritability | 0.044 | 0.053 | 0.064 | 0.011 |

**Supplementary Table 4:** Binomial logistic regression predicting remission to open-label 10 Hz and iTBS, over either the dorsomedial or dorsolateral prefrontal cortex. Greater canonical variate 2 score was associated with remission, irrespective of stimulation site or stimulation parameter. There was also a significant site * canonical variate 1 interaction, such that individuals who remitted following 10 Hz stimulation had lower canonical variate 1 scores than nonremitters, and individuals who remitted following iTBS had higher canonical variate 1 scores than nonremitters.

| **Overall Model** | **AIC** | **$R^2_{McF}$** | **$\chi^2$** | **df** | **p** |
|---|---|---|---|---|---|
| | 246.074 | 0.117 | 29.722 | 10.000 | 0.001 |

| **Predictor** | **Estimate** | **SE** | **Z** | **p** | **OR** |
|---|---|---|---|---|---|
| Intercept | -2.19 | 0.82 | -2.66 | 0.01 | 0.11 |
| Age | 0.02 | 0.01 | 1.42 | 0.16 | 1.02 |
| Sex | 0.94 | 0.38 | 2.50 | 0.01 | 2.57 |
| DLPFC > DMPFC | 0.12 | 0.62 | 0.19 | 0.85 | 1.12 |
| TBS > 10Hz | -0.28 | 0.35 | -0.80 | 0.43 | 0.76 |
| CV1 | -0.31 | 0.70 | -0.45 | 0.65 | 0.73 |
| CV2 | 1.42 | 0.57 | 2.49 | 0.01 | 4.12 |
| CV1 ✻ (DLPFC > DMPFC) | -0.36 | 0.87 | -0.41 | 0.68 | 0.70 |
| CV2 ✻ (DLPFC > DMPFC) | -0.29 | 0.76 | -0.39 | 0.70 | 0.75 |
| CV1 ✻ (TBS > 10Hz) | 1.04 | 0.51 | 2.04 | 0.04 | 2.83 |
| CV2 ✻ (TBS > 10Hz) | -0.39 | 0.71 | -0.54 | 0.59 | 0.68 |

**Supplementary Figures**

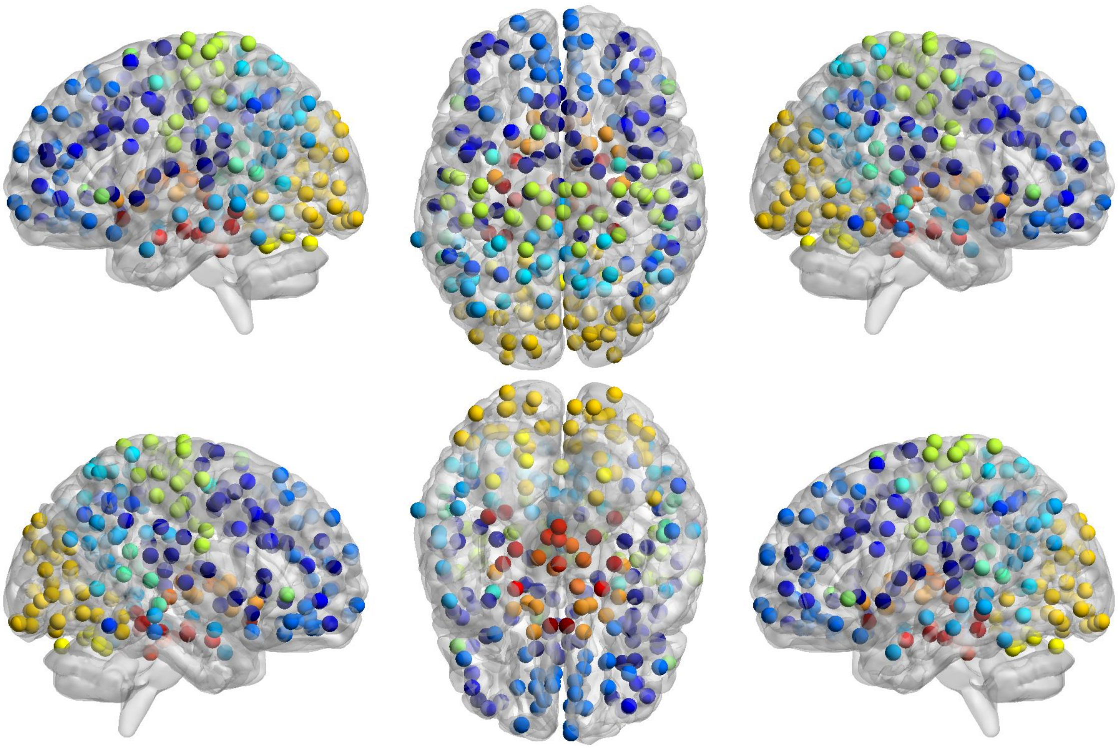

**Supplementary Figure 1:** Whole-brain RSFC was extracted from the 258 regions of interest (ROI) described in Drysdale et al.(1) resulting in 33,153 RSFC features.

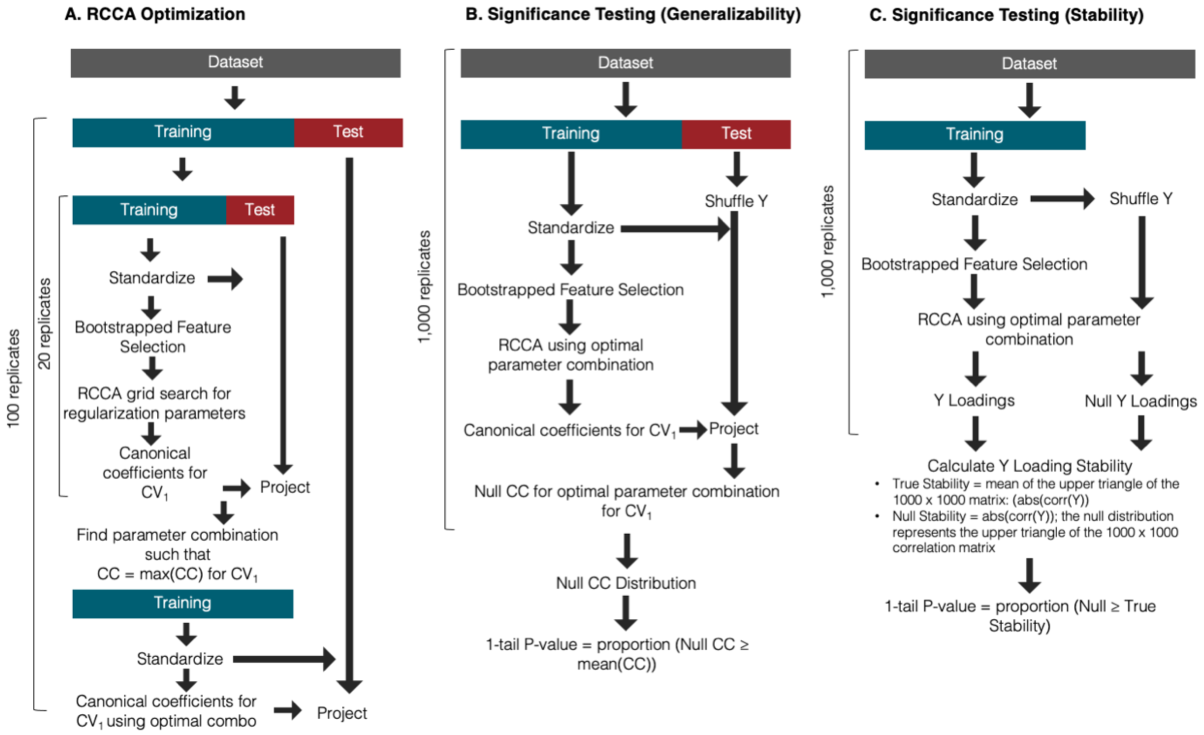

**Supplementary Figure 2:** (A) The grid search methodology used to optimize regularization hyperparameters. 219 and 109 participants were partitioned into an outer training and test set, respectively, while in the inner loop the 219 participants were further subdivided into a n = 146 and n = 73 training and test set. (B) Significance testing for assessing the performance of canonical variates in unseen data. Participants were divided into a two-thirds (n = 219) training, one-third (n = 109) test set. (C) Significance testing for assessing similarity of clinical loadings across replicates. Participants were divided into a two-thirds (n = 219) training set. Matrix X represents the RSFC matrix, where rows = participants and columns = RSFC features; Y represents the clinical matrix, where rows = participants and columns = clinical item severity.

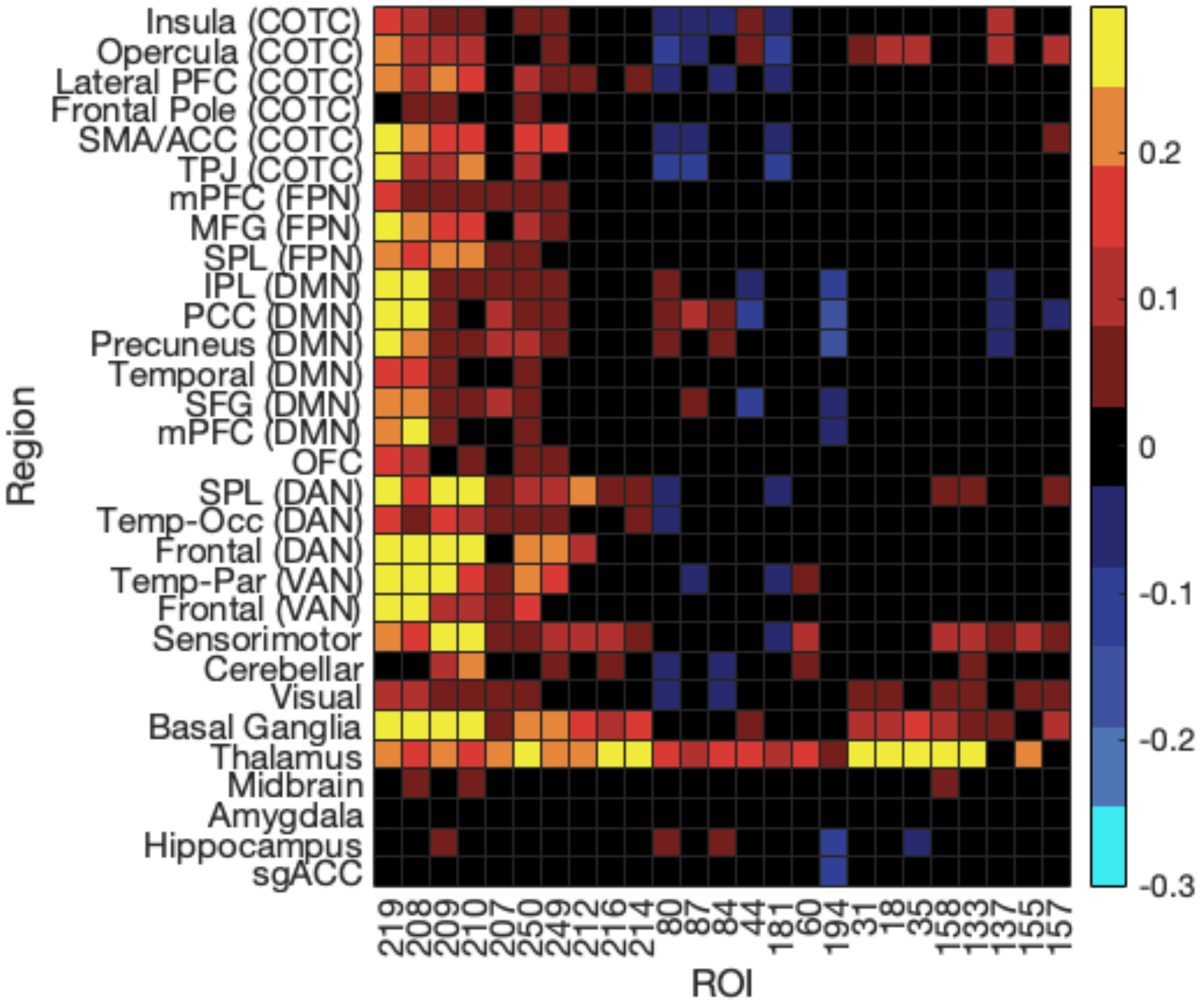

**Supplemental Figure 3:** RSFC loadings for canonical variate 1. The x-axis depicts the top 25 regions of interest, and the y-axis depicts binned regions for networks with significant features. The colorbar represents the mean Pearson correlation coefficient for features that survived FDR-correction at $p < 0.05$, whole-brain.

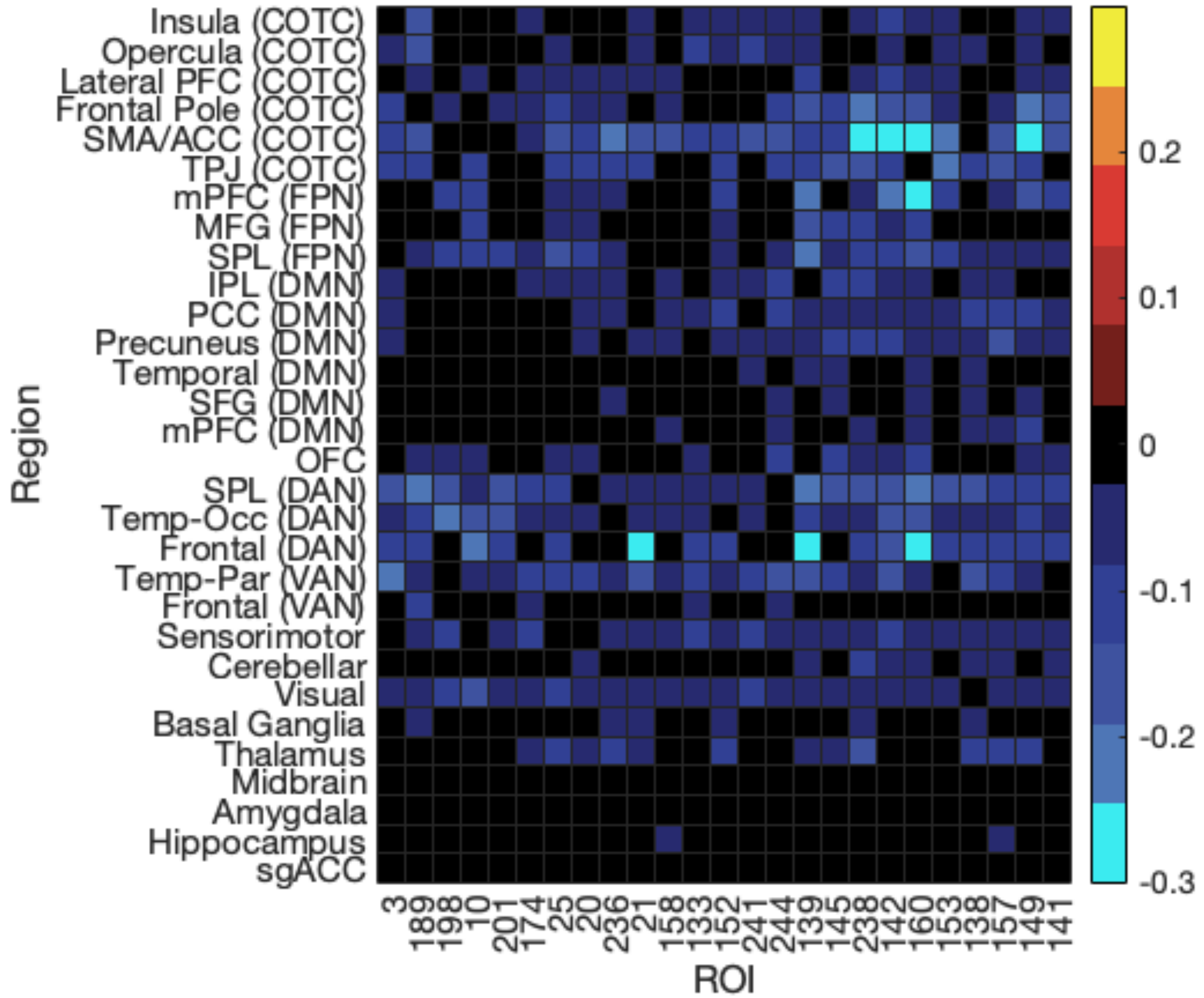

**Supplemental Figure 4:** RSFC loadings for canonical variate 2. The x-axis depicts the top 25 regions of interest, and the y-axis depicts binned regions for networks with significant features. The colorbar represents the mean Pearson correlation coefficient for features that survived FDR-correction at $p < 0.05$, whole-brain.

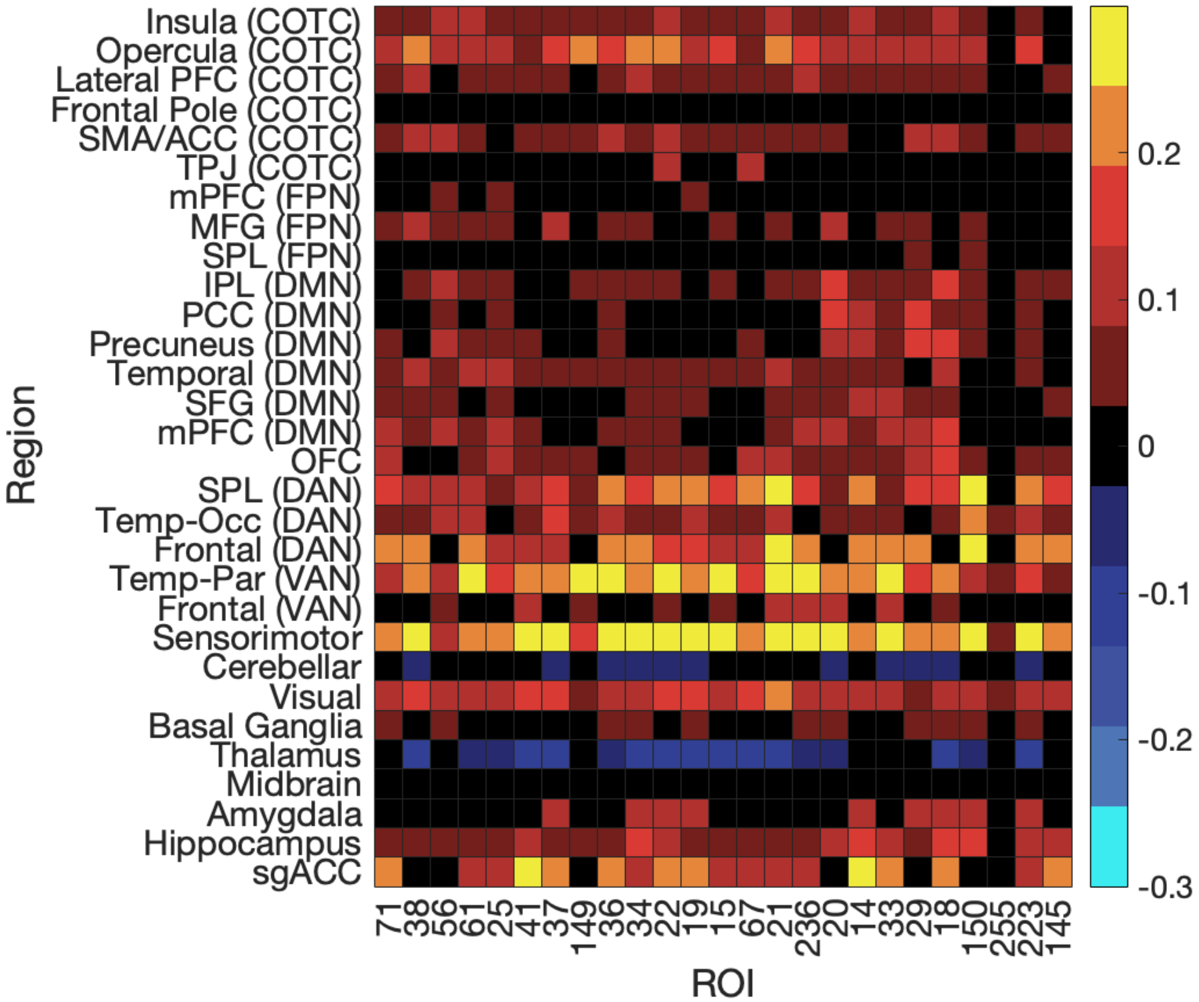

**Supplemental Figure 5:** RSFC loadings for canonical variate 3. The x-axis depicts the top 25 regions of interest, and the y-axis depicts binned regions for networks with significant features. The colorbar represents the mean Pearson correlation coefficient for features that survived FDR-correction at $p < 0.05$, whole-brain.

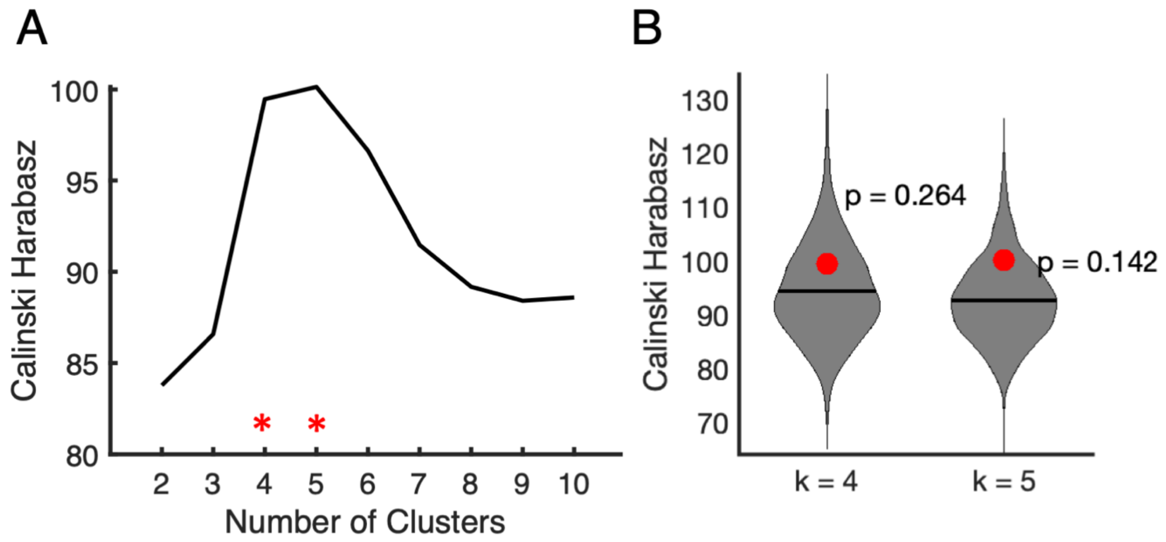

**Supplemental Figure 6:** Hierarchical clustering using all three canonical variates. (A) Calinski-Harabasz criteria values evaluating 2-10 cluster solutions when suing the first three canonical variates. Higher values indicate better performance in terms of the within- and between-cluster variance. The red asterisks indicate candidate hierarchical clustering solutions, which represents a peak in the Calinski-Harabasz criterio. (B) Permutation testing indicates that neither the $k = 4$ nor the $k = 5$ clustering solutions performs significantly better than that expected by chance. The violin plot represents the distribution of Calinski-Harabasz Criterion values for permuted training data; the red point indicates the criterion value for the unshuffled cluster solution.
